# A fungal transcription factor *BOT6* facilitates the transition of a beneficial root fungus into an adapted anthracnose pathogen

**DOI:** 10.1101/2024.03.10.584252

**Authors:** Ren Ujimatsu, Junya Takino, Masami Nakamura, Hiromi Haba, Atsushi Minami, Kei Hiruma

## Abstract

The infection strategies employed by plant endophytes are attributed to their ability to overcome durable nonhost resistance and adapt to the host environment. However, the regulatory genetic background underlying how they adapt to the host and determine their lifestyles remains enigmatic. Here, we show that the *CtBOT6*, a cluster-residing transcription factor in the root-associated fungus *Colletotrichum tofieldiae* (Ct), plays a pivotal role in regulating virulence-related gene expression and in producing metabolites both not only within and but unexpectedly outside of the cluster. Genetic manipulation of *CtBOT6* toward activation alone is sufficient to transition a root beneficial Ct along the mutualist-pathogen continuum even toward a leaf pathogen capable of overcoming nonhost resistance, partly dependent on the host abscisic acid and ethylene pathways. Our findings indicate that the status of CtBOT6 serves as a critical determinant for the endophytic fungus to adapt to the plant different environments and manifest diverse infection strategies.

## Main

Plants in nature interact with a rich diversity of endophytes that colonize inside plant tissues without causing visible disease symptoms for at least part of their life cycles^1^. Although it is little known how these endophytes can colonize inside plant tissues, they must overcome a durable and sophisticated defense system called nonhost resistance^2, 3, 4^. Even after they successfully enter host tissues, they simultaneously adapt their infection strategies, ranging from mutualistic to pathogenic, under changing host environments (changes in nutrients, host gene expression, or microbiota)^5, 6, 7, 8, 9^. Several reports have documented that host immune responses can drastically regulate the lifestyle of adapted microbes toward beneficial ones by restricting microbial colonization^10, 11, 12, 13^. From the perspective of microbes, virulence factors driving microbes from mutualistic toward pathogenic lifestyles have been characterized; the presence or absence of specific virulence factors in their genome defines pathogenicity^14, 15, 16^. Further, the activation of virulence determinants in particular fungal strains within the same fungal species or under specific environmental conditions induces a transition in lifestyle from mutualistic to pathogenic^17^. These findings suggest that microbes capable of conquering nonhost resistance can continuously define the appropriate infection strategies. However, the molecular backgrounds governing the entire infection process, from entry trial to lifestyle selection, remain elusive.

The virulence of plant-associated fungi is mediated through infection-related morphogenesis such as penetration of plant cuticles via melanized-appressoria, secretion of effectors, carbohydrate-active enzymes (CAZymes), or secondary metabolites (SM)^18, 19, 20, 21^. Transcription factors (TFs) are known to regulate the gene expression of virulence factors, and several TFs have been identified in phytopathogenic fungi^22^. Among them, Zn_2_Cys_6_ TFs are the largest TF family in phytopathogenic ascomycetes and are known not only as cluster-specific but also as global regulators of primary and secondary metabolisms^23, 24, 25^. For SM biosynthesis of phytopathogenic fungi, there is a Zn_2_Cys_6_ type TF inside the *B. cinerea* BOT cluster and the TF named BcBOT6 positively regulates the BOT gene cluster^26^. Thus far, the TF-mediated regulatory mechanisms governing the expression of virulence factors focusing on a pathogenic lifestyle have been gradually studied. On the other hand, the TF-mediated regulatory mechanisms underlying the transition in lifestyle from pathogen to mutualist or vice versa remain unexplored.

We recently reported the discovery of two strains of the root-associated endophytic fungus *Colletotrichum tofieldiae* (Ct), named Ct3 or Ct4, indicating pathogenic (plant growth inhibiting) or beneficial (plant growth promoting) lifestyles to the host *Arabidopsis thaliana*, respectively. The opposite lifestyle is attributed to a gene cluster, designated as a putative ABA-BOT cluster^17^. This cluster shares high sequence similarity with the abscisic acid (ABA) biosynthesis gene cluster and botrydial (BOT) biosynthesis gene cluster originally found in gray mold leaf pathogen *Botrytis cinerea*^27, 28, 29^ and is responsible for the biosynthesis of the intermediate metabolites of botrydial. While the putative ABA-BOT cluster exhibits little expression during root colonization by beneficial Ct4, its expression is markedly elevated in pathogenic Ct3^17^. Fluctuations in ABA-BOT gene expression are observed even within the same host species depending on environmental cues such as temperature^17^, suggesting that the expression of this cluster contributes to the switching of Ct infection strategies, ranging from pathogenic to mutualistic. However, the regulatory mechanism of the ABA-BOT gene cluster is still enigmatic. Notably, the putative ABA-BOT gene cluster also harbors a Zn_2_Cys_6_ TF within the cluster designated as *CtBOT6* in both Ct4 and Ct3, showing high amino acid sequence similarity with BcBOT6. Hence, we hypothesized that CtBOT6 regulates the ABA-BOT cluster, leading to control of the virulence to *A. thaliana*.

Host responses largely influence fungal colonization and the intensity of virulence. Plant hormones play a central role in transferring the defense signals^2, 30^. Complex regulatory mechanisms such as crosstalk between hormones result in enhanced resistance or susceptibility to plant-associated microbes^30^. While salicylic acid (SA), and ethylene (ET) have extensively been studied in the context of plant defense signaling, ABA is also important for plant-microbe interactions^31^. For instance, plant ABA signaling is required for the establishment of symbiotic interactions^32, 33^. On the other hand, in pathogenic interactions, plant ABA response contributes to susceptibility and vice versa^34, 35, 36, 37^. Although pathogenic fungi also produce ABA, its role during plant-microbe interactions has been unexplored in detail^29, 38^. While Ct activates the ABA pathway in plants in a putative ABA-BOT cluster-dependent manner, which leads to plant growth inhibition^17^, it is unclear how the ABA signaling pathway is integrated into either beneficial or pathogenic lifestyles of Ct.

In the present study, we manipulate the expression dynamics of *CtBOT6* in beneficial Ct4. Using Ct WT strains and multiple fungal transgenic lines expressing *CtBOT6* across a spectrum of expression levels, ranging from minimal to high, we observe a transition in the lifestyles of Ct4 from beneficial to pathogenic in the host plant *A. thaliana*. This transition exhibits a strong correlation with the expression levels of *CtBOT6* and, to some extent, depending on the host ABA and ET signaling pathway. Remarkably, a line with constitutive expression of *CtBOT6* can even penetrate and extensively colonize *A. thaliana* leaves, which Ct originally failed to adapt to, using melanized appressoria. This indicates that the activation of the TF is adequate to overcome nonhost resistance. Our results further suggest that the constitutive expression of *CtBOT6* leads to enhanced production of not only the botrydial intermediate metabolites but also other unknown metabolites, consistent with globally enhanced and/or diminished gene expression. These findings shed light on the central role of *CtBOT6* in fungal lifestyle transition along the pathogenic-mutualistic continuum.

## Results

### *CtBOT6* regulates the expression of the putative ABA-BOT cluster

Previously, our transcriptomic analysis revealed that Ct3 expresses a gene cluster containing ABA and BOT biosynthesis genes whereas the genes are silenced in beneficial Ct4. The gene cluster (putative ABA-BOT cluster) is composed of 10 genes, designated *CtABA1-CtABA3*, and *CtBOT1-CtBOT7*. Among the genes, we found that *CtBOT6* has a nuclear localization signal and Zn_2_Cys_6_ fungal-type DNA-binding domain, corresponding to the Zn_2_Cys_6_ TF **(Fig. 1A)**. Since *CtBOT6* is located in the putative ABA-BOT cluster, we first tried to investigate whether CtBOT6 is a TF regulating the putative ABA-BOT cluster.

**Figure 1.**
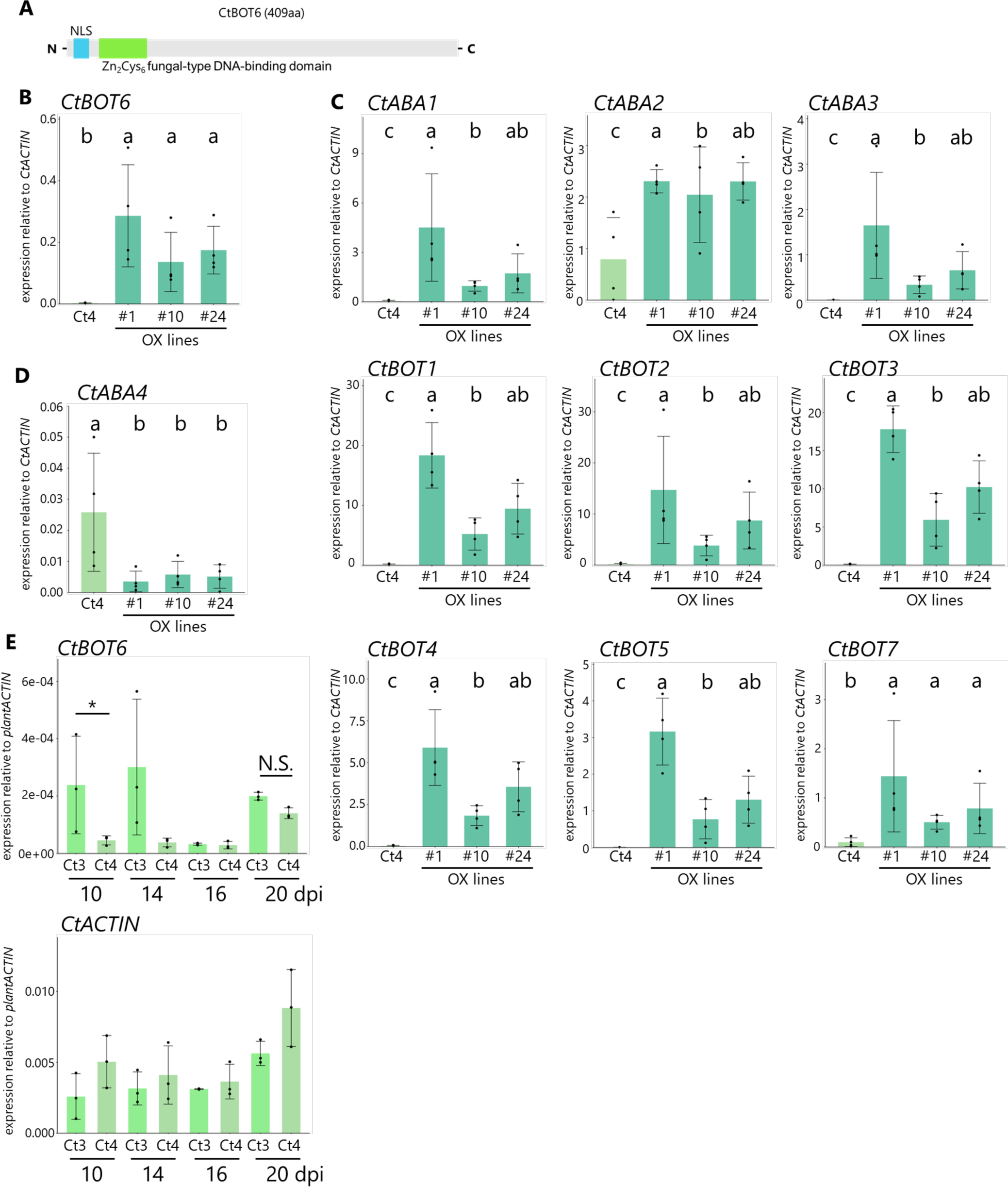
*CtBOT6* regulates ABA-BOT gene expression (A) The predicted domains of CtBOT6. The Nuclear Localization Signal was predicted using WoLF PSORT (https://wolfpsort.hgc.jp/) and Zn(II)_2_Cys_6_ DNA-binding domain (IPR001138) was predicted using Interproscan (https://www.ebi.ac.uk/interpro/result/InterProScan/). (B) *CtBOT6* expression with Ct4 WT and *BOT6* OX lines *in vitro.* Error bars represent ±SD. *n* = 4 biologically independent samples from two independent experiments. Results from two technical replicates were combined to calculate the average value in each biological replicate. Different letters indicate significant differences (adjusted *p* < 0.05). (C) Expressions of genes within Ct putative ABA-BOT cluster with Ct4 WT and *CtBOT6* OX lines (OX#1, OX#10, or OX#24) *in vitro.* Error bars represent ±SD. *n* = 4 biologically independent samples from two independent experiments. Results from two technical replicates were combined. Different letters indicate significant differences (adjusted *p* < 0.05). (D) Expressions of *CtABA4* in Ct4 WT and *CtBOT6* OX lines (OX#1, OX#10, or OX#24) *in vitro.* Error bars represent ±SD. *n* = 4 biologically independent samples from two independent experiments. Results from two technical replicates were combined. Different letters indicate significant differences (adjusted *p* < 0.05). (E) *CtBOT6* expression and Ct fungal biomass (at 10, 14, 16, 20 dpi) relative to plant *ACTIN* during *A. thaliana* root infection. *n* = 3 biologically independent samples Results from four technical replicates were combined. The asterisk indicates a significant difference (*p* = 0.01347). N.S. means no significant difference.

To examine whether CtBOT6 regulates the putative ABA-BOT cluster, we generated Ct4 transformants that constitutively express Ct*BOT6* under the TrpC promoter^39^. We have picked up three transformants with different *CtBOT6* expression levels (hereafter OX#1, OX#10, OX#24). In comparison with Ct4, OX#1, OX#10, or OX#24 induced *CtBOT6* expression 145, 69, or 89 times on average, respectively **(Fig. 1B)**. We monitored the expression of ABA and BOT biosynthesis genes in liquid media to investigate whether *CtBOT6* constitutive expression excluding other factors such as plants is sufficient to induce ABA and BOT biosynthesis genes. The quantitative RT-PCR showed that the expression of *CtABA1-3, CtBOT1-5*, and *CtBOT7* was induced by the constitutive expression of *CtBOT6* **(Fig. 1C)**. In particular, these genes were highly upregulated in transformant OX#1 *in vitro* growth condition.

In *B. cinerea*, the ABA biosynthesis cluster consists of *BcABA1*, *BcABA2*, *BcABA3,* and *BcABA4*^29, 40^. In *C. tofieldiae*, the orthologous gene of *BcABA4* is located in another contig and is not syntonically expressed with the other ABA-BOT genes during Ct3 root colonization^17^. To investigate the range of gene regulation mediated by *CtBOT6*, we measured the expression level of *CtABA4* in Ct4 and *CtBOT6* OX lines. The results indicated that the expression of *CtABA4* is not induced by the constitutive expression of *CtBOT6* **(Fig. 1D)**. Therefore, these results support the idea that *CtABA4* does not belong to the ABA-BOT cluster.

We then asked whether *CtBOT6* expression dynamics correlate with the different lifestyles between Ct3 and Ct4 on *A. thaliana*. To quantify the fungal amount and expression level of *CtBOT6* associated with *A. thaliana* root, *CtACTIN,* and *CtBOT6* were measured. RT-qPCR results highlighted that Ct3 expressed *CtBOT6,* especially in earlier time points (10, 14 dpi) **(Fig. 1E)**. Given that Ct3 inhibits the growth of *A. thaliana*, the expression of *BOT6* during the early infection stages may be required for the pathogenic lifestyle of Ct3. At 16 dpi, the expression level of *CtBOT6* in Ct3 is relatively diminished, resembling that of Ct4. This shows that *CtBOT6* is dynamically regulated during fungal colonization. Furthermore, we found that the beneficial Ct4 expressed *CtBOT6* in the late timepoint under the tested conditions (20 dpi). These results imply that moderate *CtBOT6* activation at the late stages is also required for the lifestyle of Ct4. Together, the expression level of *CtBOT6* correlates with the activation of the ABA-BOT gene cluster and flexibly changes during colonization on roots, depending on the fungal lifestyles.

### Constitutive expression of *CtBOT6* increases the production of the biosynthetic intermediates of botrydial and other sesquiterpene metabolites

We next asked whether the constitutive expression of *CtBOT6* increases the production of compounds by the putative ABA-BOT cluster *in vitro*. In our previous study, we identified two proposed intermediates for the synthesis of botrydial, 4β-acetoxyprobotryan-9β-ol (**9**) and 4β-acetoxyprobotryane-9β,15α-diol (**10**), from Ct3^17^. In this study, we subjected metabolites from *CtBOT6* OX transformants to GC-MS analysis to test the idea above. As we expected, in rice media, the amount of the intermediates **9** and **10** are drastically increased in the *CtBOT6* OX lines compared with Ct3 and Ct4 WT strains, and pathogenic *C. incanum* (Ci) WT strains wherein the ABA-BOT has **(Extended Data Fig. 1)**. Notably, compounds **9** and **10** were also detected in *CtBOT6* OX transformants when cultivated in Mathurs nutrient liquid media, whereas these metabolites were not detected in the Ct3, Ct4 or Ci WT strains **(Fig. 2)**. These results indicate that constitutive expression of *CtBOT6* in Ct4 can indeed increase the expression of the ABA-BOT genes, resulting in the high production of **9** and **10**.

**Figure 2.**
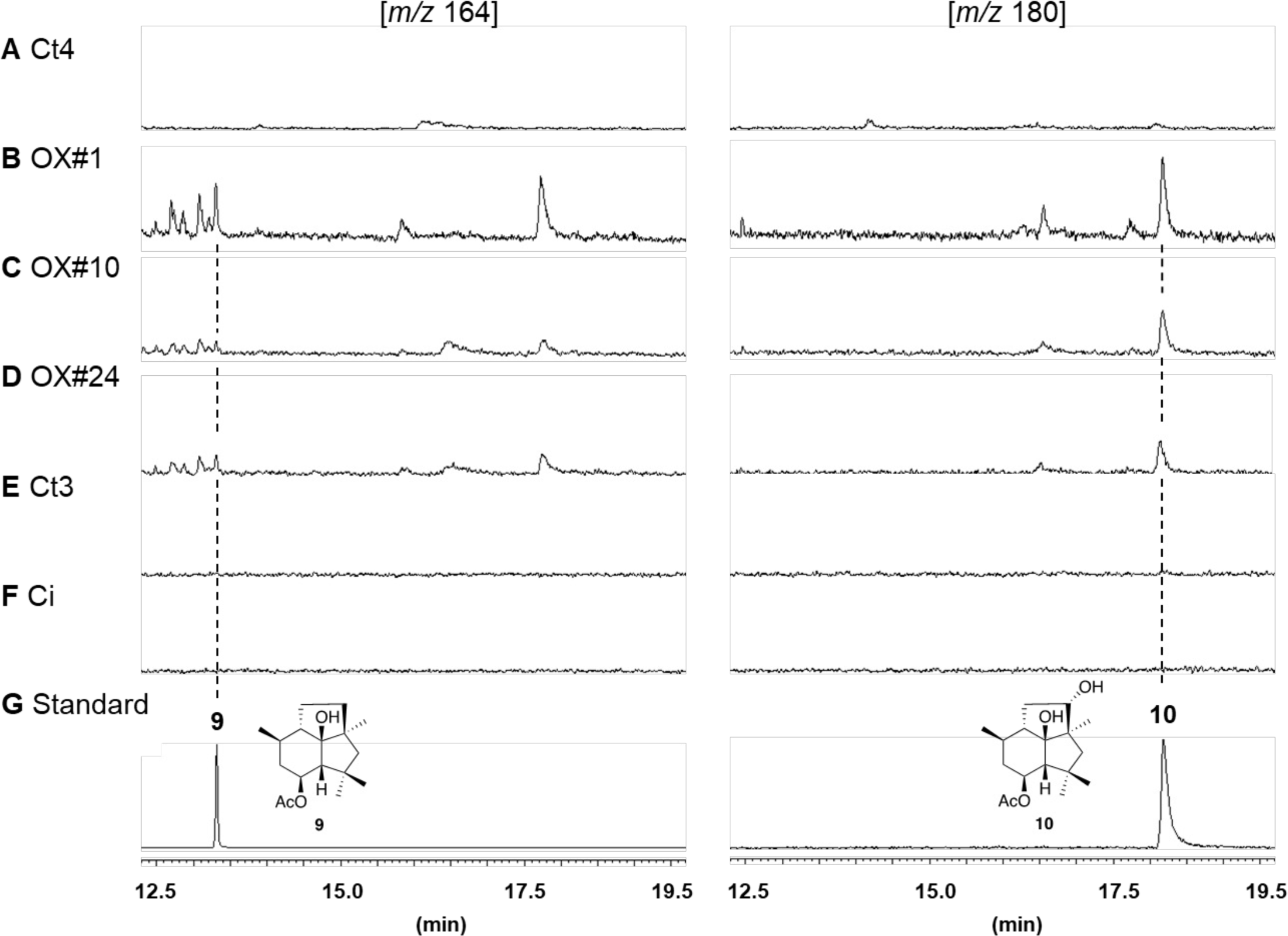
Constitutive expression of *CtBOT6* increases the production of the biosynthetic intermediates of botrydial GC-MS profiles: The metabolites (Mathurs media) from (A) Ct4, (B) OX#1, (C) OX#10, (D) OX#24, (E) Ct3, and (F) Ci. (G) standard compounds.

Next, we investigated whether Ct can also produce ABA, and/or their biosynthetic intermediates when *CtBOT6* is overexpressed. However, we were not able to detect ABA or the proposed biosynthetic intermediates when Ct transformants were grown in rice media **(Extended Data Fig. 2)**. Given the significant amino acid sequence similarity to *B. cinerea ABA* genes and the essential roles of *CtABA* genes in eliciting host ABA responses, these results imply that *CtABA* biosynthetic genes are implicated in the biosynthesis of structurally related metabolites to ABA capable of inducing host ABA responses. Remarkably, we noticed multiple peaks other than ABA or the ABA precursors in the GC-MS results, and among them, S2 was predicted to be caryophyllene, a bicyclical sesquiterpene compound, based on the similarity analysis **(Extended Data Fig. 3)**. These results imply that the constitutive expression of *BOT6* alters the dynamism of metabolite production even in vegetative growth stages in liquid media.

### Constitutive expression of *CtBOT6* alters certain aspects of fungal growth *in vitro*

Since the overproduction of metabolites originating from fungal secondary metabolism gene clusters during unintended timing is expected to consume significant energy sources^41^, and could negatively impact fungal growth, we next investigated whether constitutive expression of *CtBOT6* influences fungal vegetative growth in the absence of plants. We noticed that the transformants showed significantly reduced colony diameters *in vitro* compared to those of Ct4 and Ct3 **(Fig. 3A, B)**. The tendency is obvious in OX#1 with the highest *CtBOT6* expression **(Fig. 1B)**, implying a correlation between the expression levels of *CtBOT6* and the fungal growth. On the other hand, the number of conidia in the transformants is comparable to that of Ct4 WT, suggesting that the effects of *CtBOT6* constitutive expression are limited to specific fungal growth stages **(Fig. 3C).** Notably, the colony color of the transformants shifted to an orangish hue **(Fig. 3A)**. Given that the gene disruptions of *CtABA3* or *CtBOT1* in Ct3 showed no visible difference in colony size and color compared to the WT **(Extended Data Fig. 4)**, it suggests that the morphological and color changes may not be linked to compounds produced by the ABA-BOT gene cluster. This implies a new metabolite(s) produced by *CtBOT6* overexpression. Alternatively, ABA-BOT genes are potentially silenced or very weakly expressed in vegetative growth because of the negative effects on growth. These results indicate that the constitutive expression of *CtBOT6* had discernible effects on fungal developmental growth and metabolite production *in vitro*.

**Figure 3.**
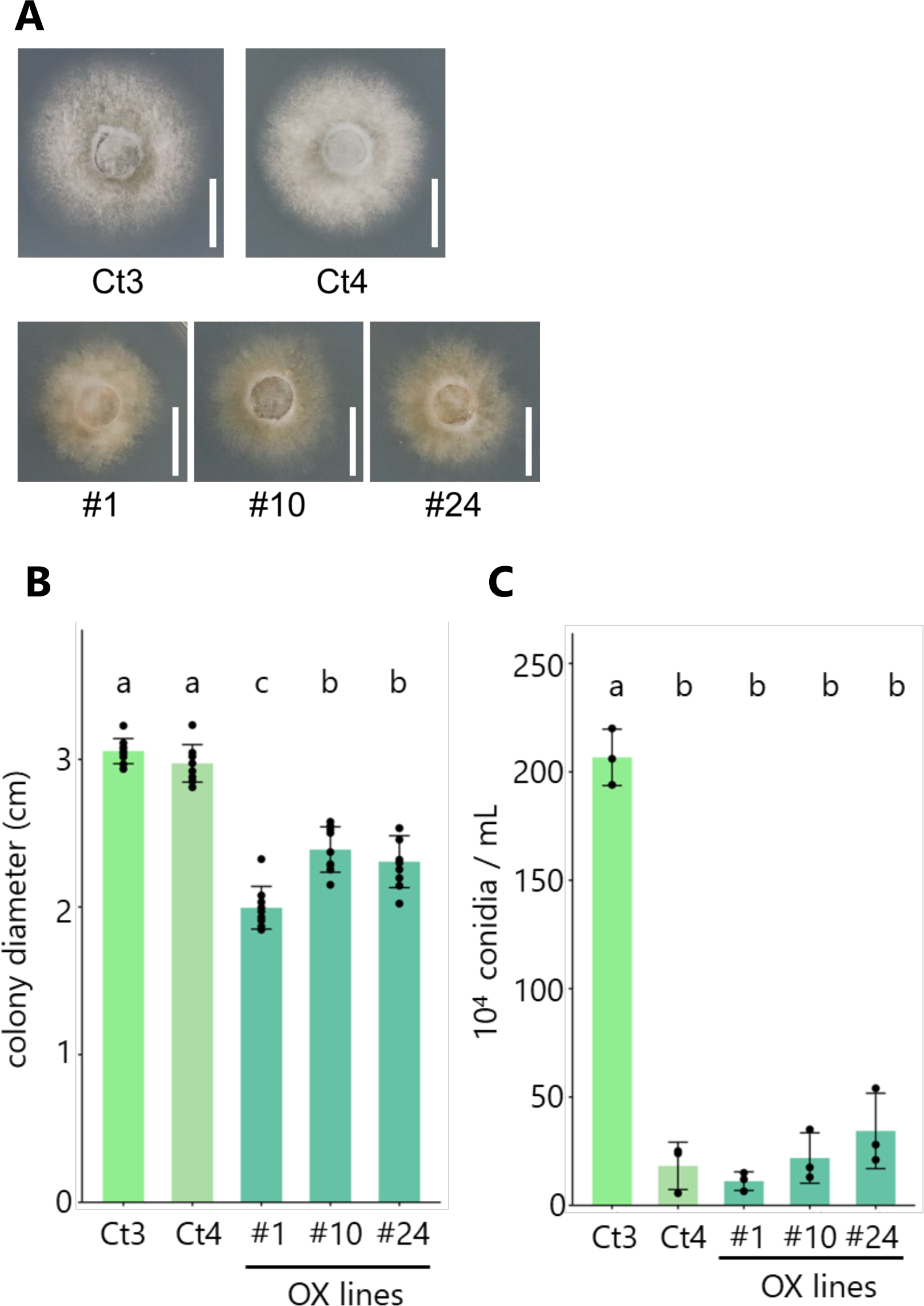
Constitutive expression of *CtBOT6* alters fungal vegetative growth *in vitro* (A) Growth phenotypes of Ct WT (Ct3 or Ct4) and *CtBOT6* OX transformants (OX#1, OX#10, or OX#24) in Mathurs nutrient media after 3 days of incubation. Bars = 1 cm. (B) Measurements of colony diameters of each fungus (Ct3, Ct4, OX#1, OX#10, or OX#24) grown in Mathurs media as indicated in Figure 3A. Different letters indicate significant differences in Tukey HSD test (*p* < 0.05). Error bars represent ±SD (*n* = 8 or 9 biologically independent samples). (C) Measurements of fungal conidia formation of Ct WT (Ct3 or Ct4) and *CtBOT6* OX transformants (OX#1, OX#10, or OX#24) grown in Mathurs nutrient media. Error bars represent ±SD. *n* = 3 biologically independent samples from three independent experiments. Different letters indicate significant differences in Tukey HSD test (*p* < 0.05).

### *CtBOT6* influences a broad spectrum of fungal secondary metabolism and virulence-related genes, alongside its impact on ABA-BOT

Our results imply that *CtBOT6* is involved in gene regulation other than ABA-BOT. To investigate the whole fungal transcriptional changes induced by *CtBOT6*, we performed RNA-seq analysis on *CtBOT6* constitutive expression transformants (OX#1, OX#10, OX#24) compared to Ct4 WT. To obtain candidate genes regulated by *CtBOT6*, we employed in vitro growth conditions in which we can exclude any potential influence from plants. Three-day-old mycelia cultured in liquid media were used for RNAseq analysis. The results revealed that 1414, 419, or 747 Ct genes were upregulated (| log2FC | >1, q < 0.05) in OX#1, OX#10, or OX#24 compared to Ct4, respectively **(Fig. 4A, Supplementary Table 1)**. The number of the upregulated gene corresponds to the expression level of *CtBOT6* in each transformant **(Fig. 1B)**. Among 14577 predicted genes of Ct4 WT, 1485 genes **(Supplementary Table 2**) or 1225 genes were upregulated or downregulated in either of OX lines **(Fig. 4A)**, respectively. To first investigate whether the 1485 up-regulated genes share motifs in the promoter regions, we conducted motif analysis on promoter regions (upper 500 bp) of 1485 genes. The MEME program predicted two significantly enriched motifs in the promoter sequences **(Fig. 4B)**. In particular, the first upper one is predicted as a highly significantly shared motif (E-value = 7.6e-025) that is shared with 564 sites in the target sequences. Our gene ontology (GO) enrichment analysis on the 1485 upregulated genes showed that GOs related to fungal metabolism (biosynthesis or metabolic processes) were highly enriched **(Fig. 4C)**, suggesting that Ct4 activates various genes related to metabolism by *CtBOT6* constitutive expression. To address this further, we predicted SM biosynthesis gene clusters and genes located in the clusters. Among the 1485 upregulated genes, 119 genes were predicted to belong to SM gene clusters **(Supplementary Table 3)**, which also supports the idea that *CtBOT6* constitutive expression up-regulates several SM gene clusters in Ct4. To address whether *CtBOT6* activation influences genes other than fungal metabolism, we next mainly focused on virulence-related genes encoding candidate secreted proteinous effectors or CAZymes regulated by *CtBOT6*. As a result, 45 of 1485 genes were predicted to be effector genes with signal peptide sequences **(Supplementary Table 4)**. 94 of 1485 genes were predicted to be CAZymes **(Supplementary Table 5)**. In summary, we obtained an unexpectedly large number of genes in OX lines differentially up-regulated genes compared to Ct4 WT, which include numerous virulence-related genes.

**Figure 4.**
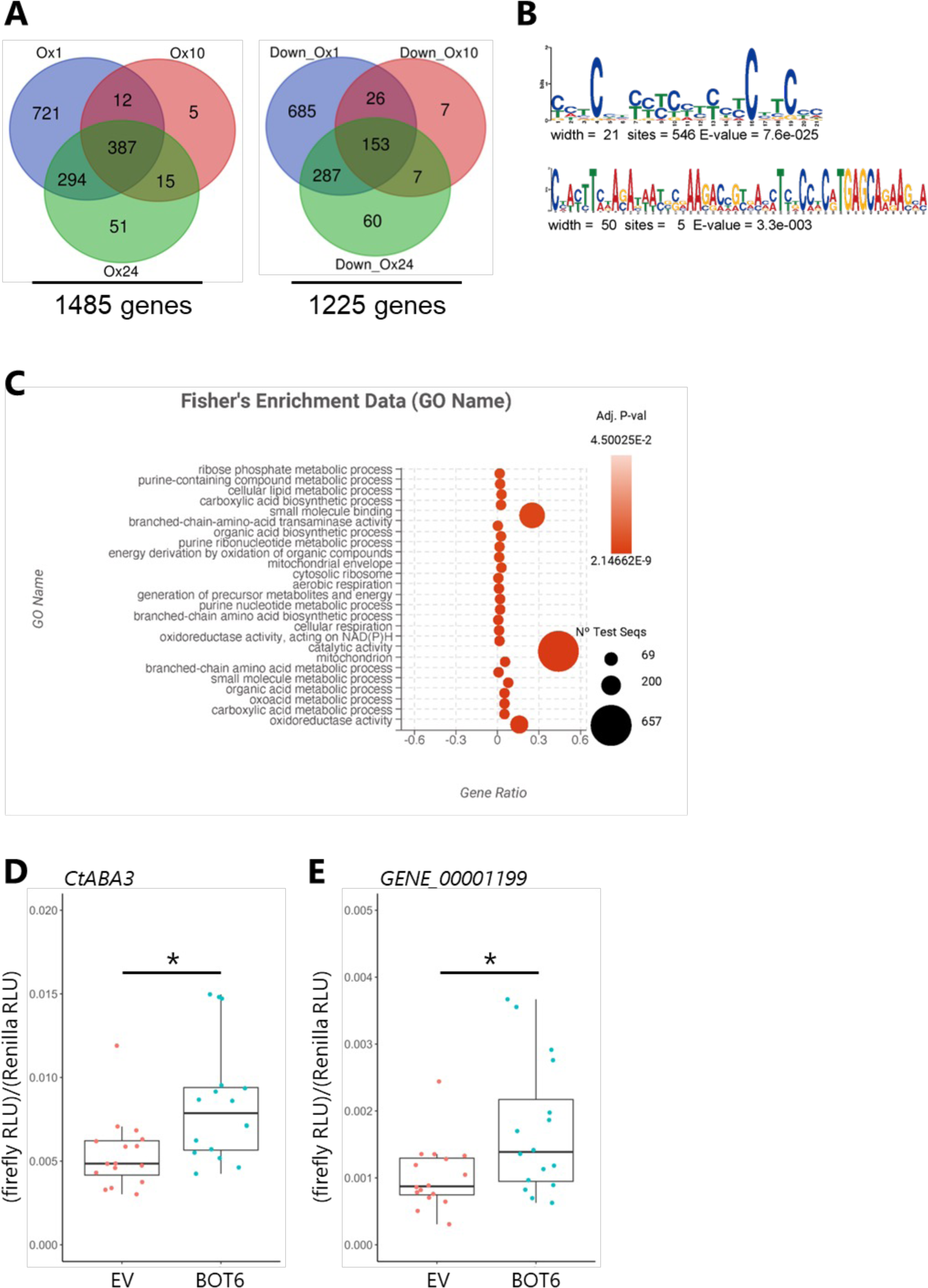
*CtBOT6* activation influences a broad spectrum of fungal secondary metabolism and virulence-related genes (A) Venn diagrams of differentially upregulated and downregulated Ct genes in three *CtBOT6* OX transformants compared with the Ct4 WT (| log2FC | >1, FDR < 0.05, Supplementary Table1). (B) Motif analysis of upper 500 bp sequences of all Ct genes upregulated in OX#1, OX#10, or OX#24 (1485 genes). The promoter sequences were subjected to the MEME analysis. (C) Gene ontology analysis shows the top 25 significantly enriched GO terms in the 1485 genes. (D) Promoter-reporter assay of *CtABA3*. *n* = 16 biological replicates from two independent experiments. Each dot represents the ratio of (firefly RLU) / (Renilla RLU, control). The asterisk indicates a significant difference (Mann-Whitney U test, *p* = 0.004484). (E) Promoter-reporter assay of GENE_00001199. *n* = 16 biological replicates from two independent experiments. Each dot represents the ratio of (firefly RLU) / (Renilla RLU, control). The asterisk indicates a significant difference (Mann-Whitney U test, *p* = 0.02108).

To investigate whether CtBOT6 can directly or indirectly regulate these genes, we tried estimating the binding ability of CtBOT6 to promoter regions of the putative ABA-BOT and/or genes located outside of the cluster. We performed a reporter assay by picking up two fungal potential target genes encoding *CtABA3*, inside of the ABA-BOT cluster, or an alcohol dehydrogenase protein, outside of the cluster (GENE_00001199) from 387 commonly upregulated genes in three transformants in the RNAseq analysis. We designed a firefly luciferase reporter driven by the fungal promoter sequences (1-2 kb) and transiently co-expressed with CtBOT6 in *Nicotiana benthamiana*. As a result, a significant increase in ratio in firefly RLU (Relative Light Units): Renilla RLU (control) was detected when the promoter sequence of *CtABA3* was co-expressed with CtBOT6 compared to the promoter sequence alone **(Fig. 4D)**. Moreover, coincubation of the promoter sequences of a GENE_00001199 and CtBOT6, also showed enhanced firefly signals significantly **(Fig. 4E)**. These results suggest that CtBOT6 can bind to promoter regions inside and outside of the putative ABA-BOT cluster. Overall, our results suggest that CtBOT6 globally regulates not only the putative ABA-BOT cluster but also other genes potentially related to virulence are regulated by CtBOT6.

### Beneficial Ct4 turns into a root and leaf pathogen by heightened expression of *CtBOT6*

We next investigated the effects of *CtBOT6* constitutive expression in beneficial Ct4 on plants during infection. To quantify the effect on plant growth, *CtBOT6* OX lines were inoculated on *A. thaliana.* Unexpectedly, we found that *CtBOT6* OX lines showed even more severe growth inhibition than pathogenic Ct3 at 24 dpi, indicating that Ct4 becomes a root pathogen when *CtBOT6* is overexpressed **(Fig. 5A, B)**. We then evaluated fungal growth under co-incubation with plants under low phosphate (Pi) conditions. First, fungal growth on half MS media with Pi-limiting (50 µM) condition was measured at 10 dpi. Notably, in contrast to *in vitro* growth **(Fig. 3A, B)**, the hyphal growth area in the OX lines is largely comparable to that in Ct4 WT strains under the co-incubation with *A. thaliana* **(Fig. 5C)**. Even more, the fungal area of OX#1 tends to be larger than that in the other OX lines, despite that the OX#1 showed the most severe growth reduction *in vitro* nutrient-rich conditions in the absence of plants **(Fig. 3A, B)**. These results imply that *CtBOT6* overexpression has a negative effect on fungal growth *in vitro* nutrient-rich conditions; however, the negative effect can be canceled in the presence of the host plant and/or in specific nutrient-limiting conditions.

**Figure 5.**
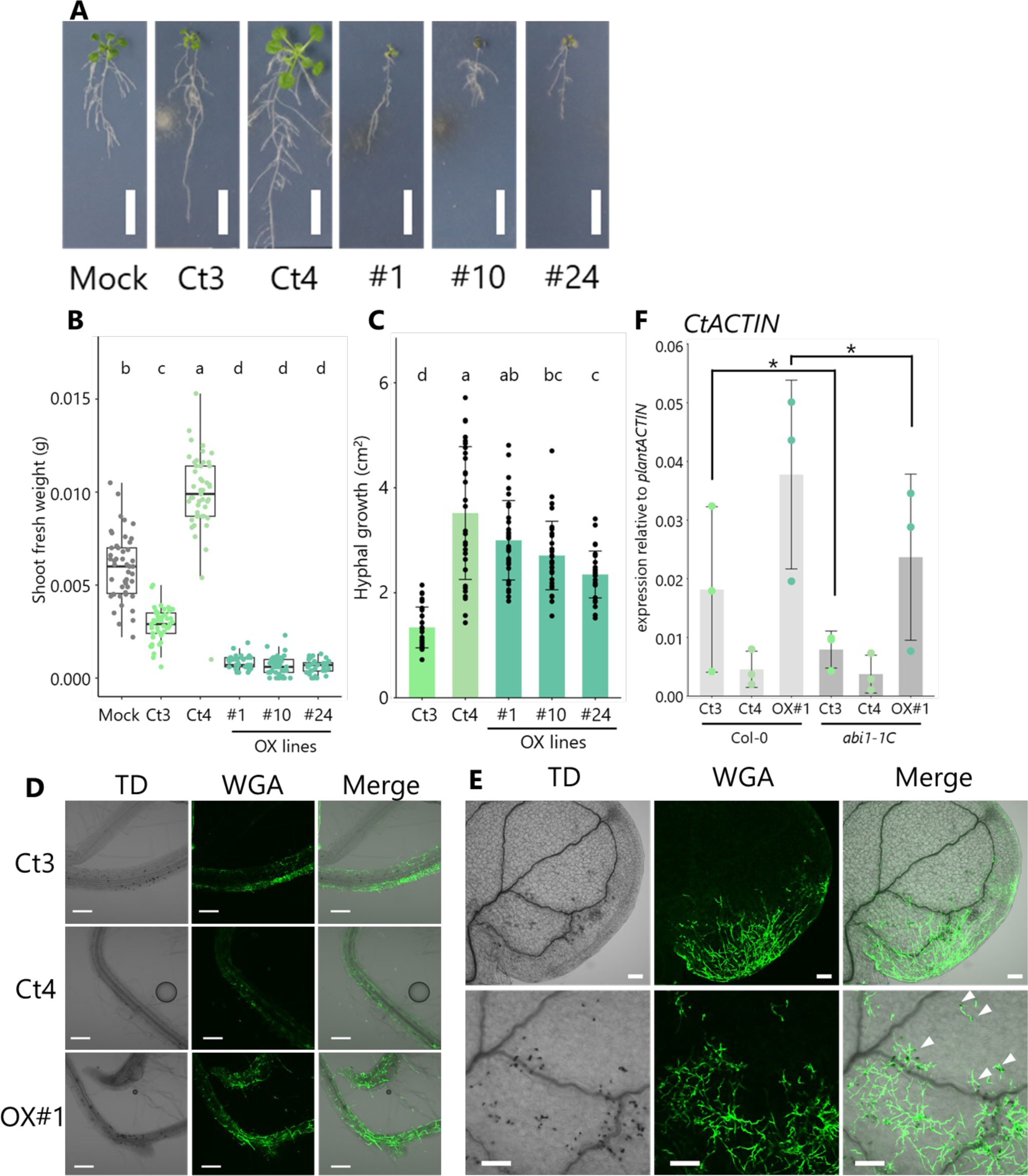
Constitutive expression of *CtBOT6* increases Ct4 root colonization and turns Ct4 into a root and leaf pathogen (A) *A. thaliana* plants grown in Pi-limiting conditions under fungal inoculation (24 dpi). Bars = 1 cm. (B) Quantitative data of *A. thaliana* shoot fresh weight in Pi-limiting conditions under fungal infection. Seedlings were harvested at 24 dpi. Boxplot represents combined results from two independent experiments. Different letters indicate significant differences in Tukey HSD test (*p* < 0.05). Each dot represents individual plant samples (Mock: *n* = 47, Ct3: *n* = 45, Ct4: *n* = 47, OX#1: *n* = 43, OX#10: *n* = 45, OX#24: *n* = 36 biologically independent samples). (C) Measurements of the fungal growth area of Ct WT and *BOT6* OX transformants grown with *A. thaliana* in Pi-limiting conditions. Error bars represent ±SD (*n* = 32 biologically independent samples from two independent experiments). Different letters indicate significant differences in Tukey HSD test (*p* < 0.05). (D) Representative confocal images of Ct hyphae stained by WGA-Lectin on *A. thaliana* root at 16 dpi (Bars = 100 μm). (E) Representative confocal images of Ct OX#1 hyphae stained by WGA-Lectin on *A. thaliana* leaves at 22 dpi (Bars = 100 μm). White arrowheads indicate invasive hyphae with melanized appressoria. (F) Fungal biomass in roots at 12 dpi. The amount of *CtACTIN* was normalized with that of plant *ACTIN*. Error bars indicate ±SD. *n* = 3 biologically independent samples from three independent experiments. Results from three technical replicates were combined. Asterisk indicates a significant difference (two tailed *t*-test, *p* = 0.02991 in Col-0 Ct3 v.s. *abi1-1C* Ct3, *p* = 0.03470 in Col-0 OX#1 v.s. *abi1-1C* OX#1).

We next observed fungal hyphae during colonization using WGA lectin staining (**Fig. 5D).** The degree of the fungal colonization on *A. thaliana* roots was obviously increased in OX#1 compared with the WT strains, further supporting the idea that Ct4 obtained severe virulence by *CtBOT6* overexpression even more than pathogenic Ct3. Notably, appressoria with melanization were frequently observed during root colonization by OX#1 as well as Ct3 compared with that by beneficial Ct4. To our knowledge, no reports are implying the role of melanized appressoria of *Colletotrichum* during root infection; however, our data implied a correlation between fungal virulence and appressoria formation on roots. Furthermore, despite that, a previous report showed that *C. tofieldiae* is not able to invade leaf tissue^10^, we even observed that OX#1 extensively colonized in *A. thaliana* leaves via direct epidermal penetration through melanized appressoria or hyphae grown from plant growth media **(Fig. 5E)**. This suggests that not only roots but also leaves are susceptible to OX#1. To further validate the over-colonization in roots by the OX line, we measured fungal biomass in roots. Consequently, there was a marked elevation in fungal biomass in OX#1 in comparison to Ct3 and Ct4 inoculation at 12 dpi **(Fig. 5F),** consistent with the microscopic observation of fungal hyphae associated with *A. thaliana* roots **(Fig. 5D)**. Interestingly, the enhanced root colonization is partially reduced in *abi1-1C A. thaliana* mutants impaired in ABA signaling whereas the fungal biomass of beneficial Ct4 was unaffected **(Fig. 5F)**. Inoculation assays revealed that the reduction of *A. thaliana* shoot fresh weights by pathogenic Ct3 and OX#1 were partially recovered in *abi1-1C*, along with the fungal biomass reduction (**Extended Data Fig. 5)**. Indeed, RT-qPCR result of *A. thaliana* ABA signaling marker gene (*MAPKKK18*) showed that host ABA signaling is enhanced in pathogenic Ct3-infected roots and even higher but not significantly upregulated in *CtBOT6* OX#1 in comparison with Ct4 **(Extended Data Fig. 6)**. These results suggest that the constitutive expression of *CtBOT6* markedly enhanced the hidden virulence of the beneficial Ct4 strain, partially by eliciting the activation of host ABA signaling, during colonization of *A. thaliana* roots.

### Plants enhance defense responses along with the fungal virulence independent of the ABA signaling pathway

Plants mount defense responses to counteract microbial infections. A beneficial strain of Ct (Ct61) still can promote plant growth in *A. thaliana* mutant disrupted in SA, JA, ET, and *AtPAD4*^10^. However, the potential effects on plant defense response by pathogenic Ct invasion have been unexplored. Additionally, it is currently unknown whether there is crosstalk between the plant ABA signaling pathway and other plant defense responses in the Ct-*A. thaliana* interaction. To investigate these questions, we utilized marker genes of SA (*PBS3*), ET (*ETR2* and *ERF1*), tryptophan-derived antifungal pathway (*CYP71A12* and *CYP81F2*), and *FRK1*, a marker gene for PAMP (pathogen-associated molecular pattern)-triggered immunity (PTI). The expression levels of these genes are significantly higher in Ct3 compared with Ct4 **(Fig. 6)**. Besides, even stronger gene expression is detected in OX#1 for ET marker genes, tryptophan-derived antifungal pathway marker genes, and *FRK1* **(Fig. 6)**. These results indicate that activation of *CtBOT6* in Ct upon the fungal invasion heightened defense responses, notwithstanding the increased fungal biomass **(Fig. 5F)**. Notably, although the ABA signaling pathway plays a role in fungal virulence **(Fig. 5F, Extended Data Fig. 5)**, the hyperactivation of defense responses observed here are unaffected in *abi1-1C* **(Fig. 6)**. Together, it is suggested that at least some defense genes located in several different pathways are activated along with *CtBOT6* expression levels and that seems not influenced by ABA PYR/PYL-PP2C signaling core pathway.

**Figure 6.**
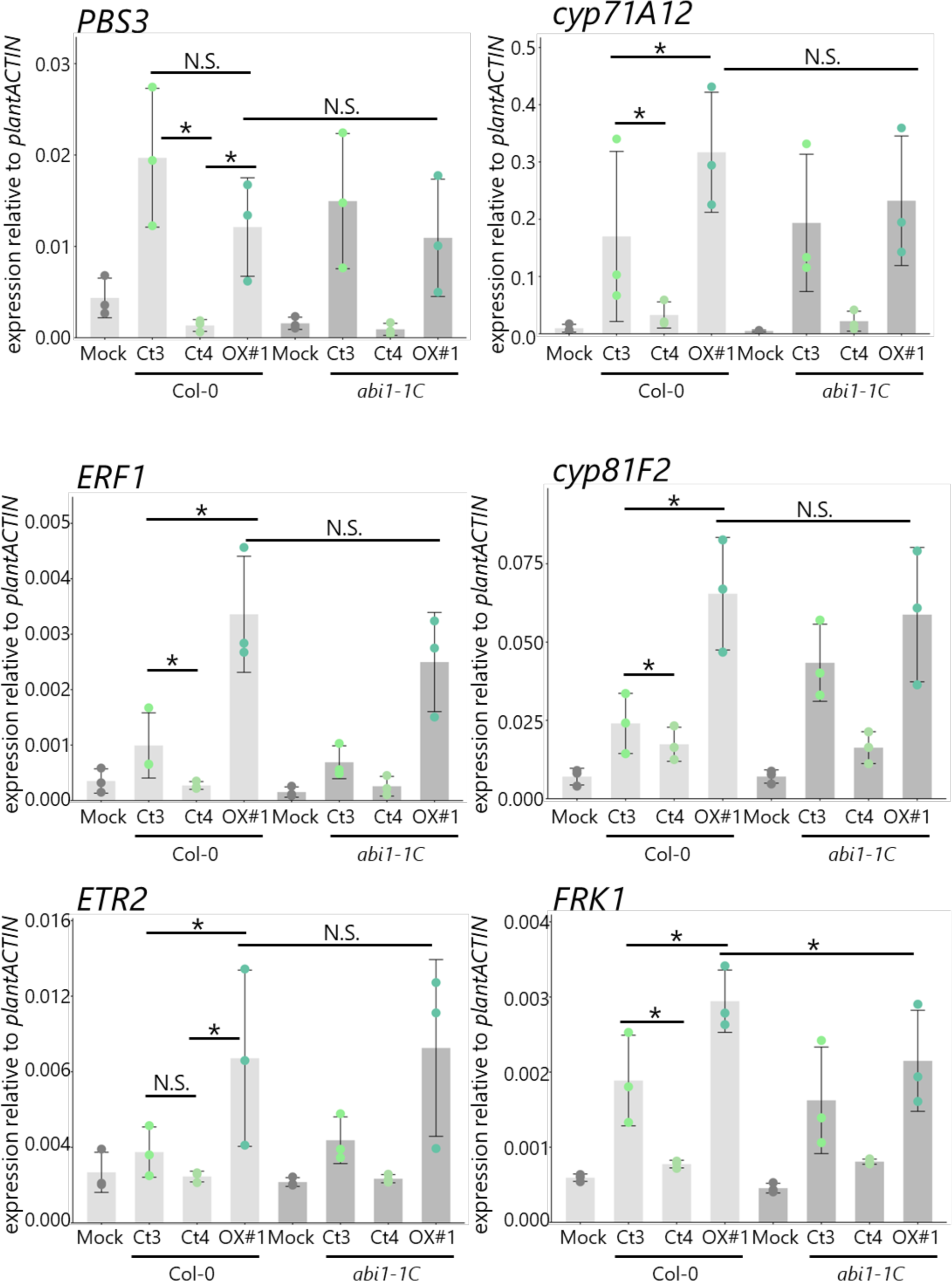
Plants activate defense-related genes in response to fungal virulence facilitated by *CtBOT6* Gene expression levels in *A. thaliana* roots at 12 dpi, normalized to the reference plant ACTIN, were measured by RT-qPCR. Error bars indicate ±SD. *n* = 3 biologically independent samples from three independent experiments. Results from three technical replicates were combined to obtain the mean from each independent experiment. The asterisks indicate a significant difference (two tailed *t*-test, *PBS3*: *p* = 0.0000029 in Col-0 Ct3 v.s. Col-0 Ct4, *p* = 0.0000283 in Col-0 Ct4 v.s. Col-0 OX#1; *ERF1*: *p* = 0.00073 in Col-0 Ct3 v.s. Col-0 Ct4, *p* = 0.00043 in Col-0 Ct3 v.s. Col-0 OX#1; *ETR2*: *p* = 0.00459 in Col-0 Ct3 v.s. Col-0 OX#1, *p* = 0.00018 in Col-0 Ct4 v.s. Col-0 OX#1; *cyp71A12*: *p* = 0.000037 in Col-0 Ct3 v.s. Col-0 Ct4, *p* = 0.00637 in Col-0 Ct3 v.s. Col-0 OX#1; *cyp81F2*: *p* = 0.03218 in Col-0 Ct3 v.s. Col-0 Ct4, *p* = 0.0000009 in Col-0 Ct3 v.s. Col-0 OX#1; *FRK1*: *p* = 0.000042 in Col-0 Ct3 v.s. Col-0 Ct4, *p* = 0.00615 in Col-0 Ct4 v.s. Col-0 OX#1, *p* = 0.03645 in Col-0 OX#1 v.s. *abi1-1C* OX#1). N.S. means no significant difference.

Since ET pathways are activated during root colonization by Ct3 and OX#1, we tested whether ET signaling is involved in the virulence caused by these fungi. Inoculation of these fungi individually with *ein3 eil1* mutants, which are defective in ET pathway, revealed a reduction in the virulence, as indicated by effect size **(Extended Data Fig. 7)**. This suggests that, in addition to ABA pathways, ET pathway is required for the virulence caused by Ct3 and OX#1.

### Genetic manipulation of *CtBOT6* expression reveals that *CtBOT6* expression level is positively correlated with the degree of virulence in Ct4

To further investigate whether the expression amount or timing of *CtBOT6* during root colonization is correlated with fungal virulence, we designed a transformant that expresses *CtBOT6* under regulation of a Ct3 phosphate transporter promoter (GENE_00003000, hereafter phos *pro*), whose gene in Ct3 and the orthologous gene (GENE_00011871) in Ct4 are highly expressed in low phosphate condition especially at the late infection stage^17^. We first monitored the expression of the phosphate transporter gene during root colonization by Ct4 under our tested condition. As a result, the expression patterns of the gene in Ct4 are consistent with the previous study^17^ **(Extended Data Fig. 8)**, suggesting that the phos *pro* is useful for expressing *CtBOT6* at varying levels, in a dependent manner of environmental Pi conditions **(Fig. 7A)**. We generated 14 lines of the transformants (denoted as phos lines), and we have picked up three of them (phos#1, phos#4, phos#6). Inoculation assay of these transformants on *A. thaliana* roots resulted in diverse outputs from mild growth inhibition to plant growth promotion **(Fig. 7B, C)**. In detail, phos#1 inhibited plant growth fairly close to Ct3 WT, whereas phos#4 promoted plant growth second to Ct4 WT. phos#6 showed mild plant growth inhibition compared to phos#1. Next, we investigated whether the variety of virulence against plants is correlated with *CtBOT6* expression level. RT-qPCR results of the roots from three independent experiments exhibited that, among the phos transformants, phos#1 showed the highest expression of *CtBOT6* especially in earlier timepoints and simultaneously induced the lowest plant shoot fresh weight **(Fig. 7C, D)**. The tendency of *CtBOT6* expression and the shoot fresh weight in phos#1 resembled pathogenic Ct3. phos#4 showed similar *CtBOT6* expression dynamics to beneficial Ct4 with a similar shoot fresh weight. phos#6 displayed a middle phenotype both in *CtBOT6* expression levels and the shoot fresh weight. Furthermore, Pearson’s product-moment correlation coefficient showed a significant correlation between shoot fresh weight and *CtBOT6* expression levels at 22 dpi (p-value = 0.003562). Importantly, a significant correlation was even detected between shoot fresh weight at 22 dpi and *CtBOT6* expression levels at 14 dpi (p-value = 0.01732) or at 18 dpi (p-value = 0.01483). These results indicate that the degree of virulence is positively correlated with the expression level of *CtBOT6*. Thus, our results indicate that Ct4 can transition lifestyles along the mutualist-pathogen continuum through *CtBOT6* expression changes.

**Figure 7.**
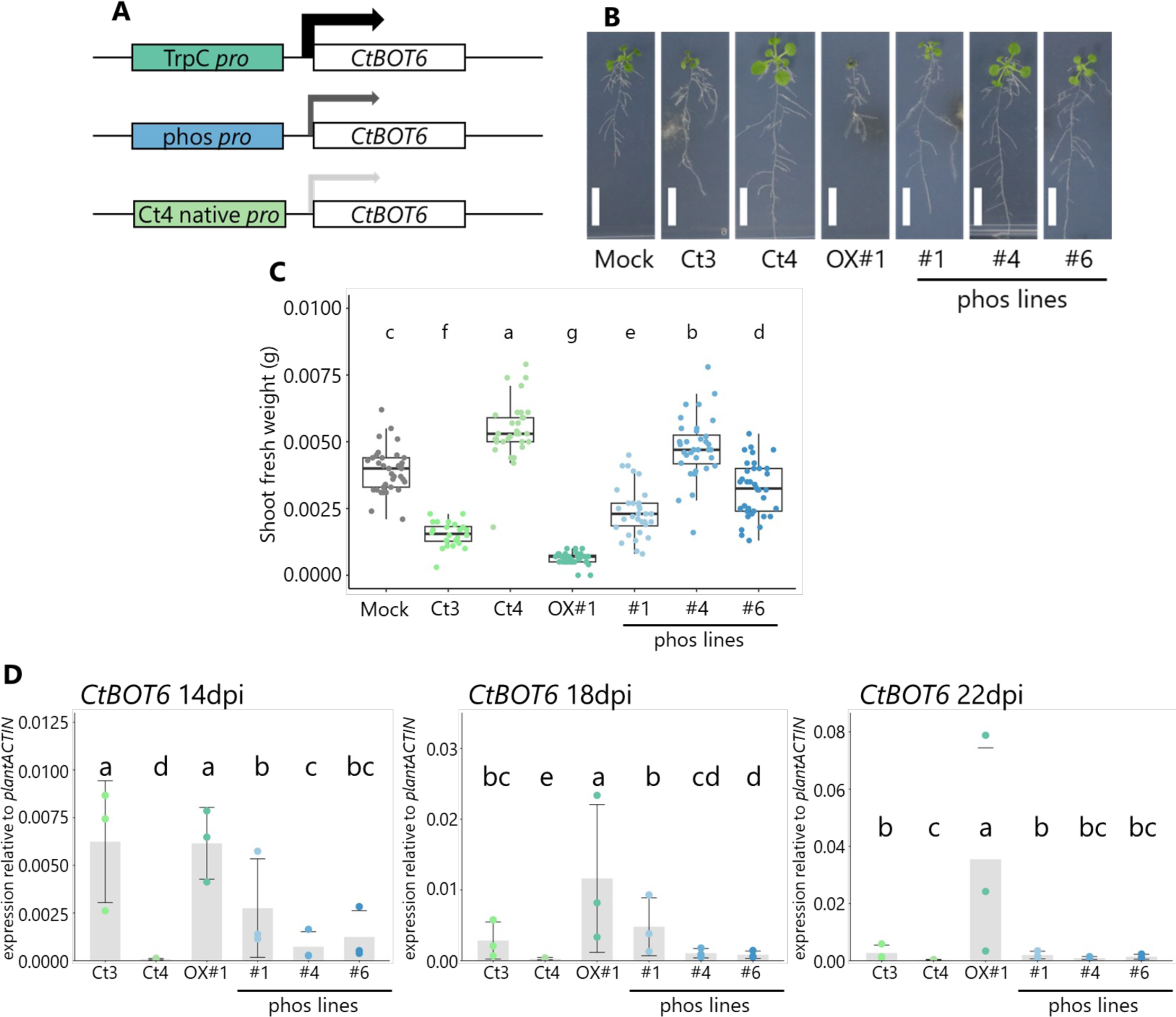
*CtBOT6* expression level is positively correlated with the degree of virulence in Ct4 (A) Structure of genome or construct designed in this study expressing CtBOT6 at varying levels. (B) *A. thaliana* plants co-incubated with the described fungi in Pi-limiting conditions (22 dpi). Bars = 1 cm. (C) Quantitative data of the shoot fresh weight on *A. thaliana* inoculated with the fungi in Pi-limiting conditions. Seedlings were harvested at 22 dpi. Different letters indicate significant differences in Tukey HSD test (*p* < 0.05). Each dot represents individual plant samples (Mock: *n* = 35, Ct3: *n* = 24, Ct4: *n* = 33, OX#1: *n* = 35, phos#1: *n* = 31, phos#4: *n* = 36, phos#6: *n* = 36 biologically independent samples). (D) *CtBOT6* expression (14, 18, 22 dpi) amount relative to plant *ACTIN* during *A. thaliana* root infection. *n* = 3 biologically independent samples from three independent experiments. Results of three technical replicates were combined to obtain the mean from each independent experiment. Error bars represent ±SD. Different letters indicate significant differences (adjusted *p* < 0.05)

## Discussion

In this study, we show that genetic manipulation of only one TF *CtBOT6* in beneficial root-associated Ct4 is sufficient to elicit the hidden virulence against *A. thaliana* **(Fig. 5)**. The virulence induced by *CtBOT6* enabled Ct4 to infect not only *A. thaliana* roots but also *A. thaliana* leaves, where Ct is naturally non-adapted^10^, through a conventional entry mode of leaf pathogenic *Colletotrichum* using black melanized appressoria **(Fig. 5E)**. These indicate that *CtBOT6* activation transforms Ct4 into not only a root pathogen but also a leaf anthracnose pathogen capable of overcoming nonhost resistance, a robust defense system comprising both pre- and post-invasion defense layers in leaves^10, 42, 43^.

It has been reported that microbial TFs regulate the expression of virulence factors and are responsible for virulence in various parasites infecting various host organisms^22^. Further, it has been described that expression levels of a bacterial *Xenorhabdus nematophila* TF Lrp determine the lifestyles on different host organisms^44, 45^. However, little is known about whether and how the expression level of a TF governing virulence determines the diverse lifestyles of a microbial species within the same host. Here, we analyzed fungal strains expressing *CtBOT6* at various levels, ranging from minimal (Ct4) to moderate (phos4 and phos6), high (Ct3 and phos1), and very high (OX#1), utilizing two independent promoter sequences in addition to the native promoter sequences of *CtBOT6* in Ct3 and Ct4 **(Figs. 5, 7)**. As a result, we discovered that the intensity of Ct virulence on *A. thaliana* strongly correlated with the expression level of the TF. Considering the diverse lifestyles among Ct strains under different environmental conditions, this result implies that Ct optimizes its intensity of virulence by fluctuating the expression of *CtBOT6*. Interestingly, there is a negative impact on vegetative growth in nutrient-rich media when *CtBOT6* is constitutively expressed; however, under the interaction with the host plant on Pi-limiting media, the OX lines can grow on media at a similar level to the WT Ct4 and infect roots excessively **(Figs. 3, 5C)**. These suggest that the effect of *CtBOT6* constitutive expression on fungal growth can be both positively and negatively affected by the environmental conditions including the host, which might explain that *CtBOT6* is regulated and is specifically detected during root colonization by pathogenic Ct3 but not beneficial Ct4^17^. Moreover, it has been reported that ABA-BOT gene expression in Ct3 becomes restricted at the later colonization phase (**Fig. 1E**)^17^. In relation to this, despite that constitutive expression of *CtBOT6* leads to a drastic increase in Ct4 biomass in *A. thaliana* roots as well as leaves, the conidia and setae formation that is typical to *Colletotrichum* pathogen are not observed in both tissues even at 22 dpi. In the context of roots, this may be attributed to the high induction of plant defense responses, some of which are known to restrict beneficial Ct growth in roots **(Fig. 6)**^10^. This may result in the suppression of fungal reproduction during late root colonization by *CtBOT6* OX lines. Therefore, these findings suggest the importance of *CtBOT6* expression at precise timing across the fungal life cycle not solely for fungal infection strategies but also for fungal reproduction.

Plant ABA signaling plays a crucial role in plant-microbe interactions. Here we show that the host ABA signaling pathway contributes to the increase of fungal biomass on plant roots **(Fig. 5F)**. ABA signaling enhances plant susceptibility toward *B. cinerea*, known as gray mold^37^. Knockout of the *WRKY33* gene, a defense-related TF, enhances ABA biosynthesis in plants in a dependent manner on *NCED3* and *NCED5* ABA biosynthesis genes. This resulted in a marked increase in fungal colonization of plant leaf tissues. Indeed, the nonadapted *B. cinerea* can hardly infect *wrky33 nced3 nced5* mutant plants^37^, indicating that activation of the host ABA pathway via WRKY33 suppression is critical for virulence. It remains unclear whether Ct with *CtBOT6* activates ABA signaling through a comparable host pathway (potentially invoking suppression of WRKY33) as *B. cinerea* presumably does. Nevertheless, given that *B. cinerea* also has ABA and BOT biosynthetic genes in its genome and activates host ABA signaling, it is plausible that Ct and *B. cinerea* have analogous strategies to activate host ABA signaling during root or leaf colonization, respectively.

On the other hand, our data also suggested that the contribution of the ABA pathway is relatively limited in Ct*-A. thaliana* interaction because the virulence and root colonization of Ct3 and *CtBOT6* OX#1 are still retained in *abi1-1C* mutant plants **(Fig. 5F)**. Given that the *abi1-1C* mutant lacks, not all, but a large part of the ABA signaling^46^, it is reasonable to consider that in addition to the ABA pathway, unknown factors contribute to the fungal virulence. Consistent with this hypothesis, the induction of SA, ET, tryptophan-derived antifungal biosynthesis, and PTI marker genes during root colonization by pathogenic Ct was unaffected in *abi1-1C* **(Fig 6)**. Furthermore, ET signaling pathway seems to be required for the fungal virulence **(Extended Data Fig. 7)**. These results suggest that other pathways such as ET are also involved in Ct virulence in addition to host ABA signaling.

From fungal side, our transcriptome analysis unveiled that *CtBOT6* constitutive expression in vitro (i.e., without plants) altered the expression of almost 20% of Ct genes including several virulence-related genes in either of three OX lines **(Fig. 4A) (Supplementary tables 4-5)**. Considering these results, it is plausible that multiple factors regulated by CtBOT6 are combined to support Ct virulence in this condition. Interestingly, our previous phylogenetic analysis on the putative ABA-BOT cluster suggests that *BOT6* is distributed in plant-associated pathogenic fungi in which *ABA* and *BOT* genes do not seem to exist, suggesting a different evolutionary background of *CtBOT6* from other ABA/BOT genes^17^. Moreover, given that there are some examples of TFs inside of a cluster but regulating outside of the cluster^47, 48^, it can be hypothesized that *CtBOT6* is responsible for the expression of not only within the cluster but also outside the cluster. This mechanism by which CtBOT6 influences various potential targets would be important for understanding the hidden regulatory mechanisms of Ct virulence along the parasitic-mutualistic continuum.

Our attempts to transform the pathogenic Ct3 into a mutualist by disrupting the *CtBOT6* gene were unsuccessful, despite our efforts to try to knockout the gene in both Ct3 and Ct4 using two independent vectors for knockout. Nevertheless, considering that the knockout of *CtBOT5* within the ABA-BOT in pathogenic Ct3 was adequate to shift its pathogenic lifestyle to a beneficial one under Pi-limiting conditions^17^, it is plausible to infer that *CtBOT6* knockout in Ct3 if generated would lead to a transition in lifestyle from pathogenic to beneficial. Conversely, unlike the other ABA and BOT genes, the inability to generate *CtBOT6* knockout fungi implies that *CtBOT6* might be indispensable for fungal survival. This is noteworthy compared with a previous report that *BcBOT6* was disrupted in *B. cinerea* without affecting the virulence and growth of the fungus^26^. It has been estimated that *BOT* genes including *CtBOT6* undergo horizontal gene transfer (HGT) from *Colletotrichum* to *Botrytis*^17^, thus our findings may imply a functional divergence between *CtBOT6* and *BcBOT6* following HGT.

In conclusion, our results suggest that the dynamic expression of *CtBOT6* contributes to the parasitic–mutualistic continuum observed in Ct infection strategies. The fluctuation of expression levels of *CtBOT6* during infection in both Ct3 and Ct4 WT implies that *CtBOT6* expression is sensitively influenced by host and environmental changes at each infection stage, which may determine the continuous infection strategy of Ct under changing environments. However, the regulatory mechanisms governing *CtBOT6* expression are still enigmatic. At least, fungal TFs are known to be influenced by environmental nutrient availability, which is hypothesized to undergo continuous fluctuations within the host environment^47, 49^. Further investigations aimed at elucidating these regulatory mechanisms in response to environmental cues will hopefully expand understanding of the mechanisms underlying the discrimination of pathogenic and beneficial fungal lifestyles, as well as the diversity and continuum of fungal infection strategies and their adaptive responses to the host environment.

## Materials & Methods

### Fungal strains, media, and transformation

The *Colletotrichum tofieldiae* Ct3 (MAFF712333) and Ct4 (MAFF712334) strains and the *Colletotrichum incanum* (MAFF238704) strain were used as the wild-type strains in this study^17^. Ct3 knockout mutants of *Ctaba3*, *Ctbot1*#27, and *Ctbot1*#30 were generated in the previous study^17^. Fungi are maintained on Mathurs media with 3% agar granulated (Difco) at 25°C in normal light conditions. Fungal transformation was performed as previously described^17^. Briefly, the knock-in vector (see below) was introduced into Agrobacterium C58C1 strains by electroporation. The transformed Agrobacterium strains (OD600 = 0.4) were mixed with Ct4 spores (1 × 10^7^) in a 1:1 ratio on a paper filter attached to incubation media containing 200 μM acetosyringone (Fuji Film). Selection of hygromycin (150 μM)-resistant strains was conducted by transferring the paper filter to PDA media (Difco) with Hyg, cefotaxime, and spectinomycin (all at 200 μg/mL) and incubated for two days before the paper was eliminated from the media. The resistant strains were transferred to new media with three antibiotics and confirmed again whether they showed resistance against Hyg.

### Cloning

*Escherichia coli* Stellar^TM^ Competent Cells (Clontech) was used as the host for DNA manipulation. All primers used in this study are listed in Supplementary Table 6. Templates used for PCR were derived from gDNA or cDNA of *C. tofieldiae* strain Ct3 and Ct4. All DNA fragments were amplified from templates by PCR reaction using KOD One PCR Master Mix (TOYOBO), PrimeSTAR® GXL DNA Polymerase (Takara), or Prime STAR HS DNA Polymerase (TaKaRa) with standard protocols.

To generate a knock-in vector for constitutively expressing *CtBOT6* in Ct4, amplified *CtBOT6* gene was substituted with the GFP gene within the binary plasmid pBin-GFP-hph^50^. The pBin-GFP-hph vector devoid of the GFP gene region was amplified. The *CtBOT6* DNA fragment and the amplified vector were then merged using the In-Fusion HD Cloning Kit (TaKaRa). Ct4 transformed with the pBin-GFP-hph-*CtBOT6* vector express *CtBOT6* under the TrpC promoter control. The successfully isolated strains were incubated in liquid media for three days and were tested by RT-qPCR after RNA extraction using primers targeting the *CtBOT6* gene sequence to determine whether the *CtBOT6* was successfully over-expressed. To generate a knock-in vector for expressing *CtBOT6* under the control of the Ct3 phosphate H^+^ symporter gene promoter in Ct4, the promoter region of Ct phosphate symporter gene (the upstream of 1843 sequences of GENE_00003000), and *CtBOT6* gene following the promoter sequences were replaced with the GFP gene, including TrpC promoter region, within the pBin-GFP-hph vector. The pBin-GFP-hph vector lacking the GFP gene region and the TrpC promoter region was amplified. Subsequently, the promoter of the phosphate symporter gene and *CtBOT6* DNA fragment were fused with the PCR amplified vector by infusion. Ct4 transformed with this vector expresses *CtBOT6* under the regulation of the promoter of the phosphate symporter gene. For dual luciferase assay, amplified promoter region fragments are inserted with KpnI and XhoI site in pAT006-Fluc-iG5_Rluc-iFAD2_F vector using In-Fusion HD Cloning Kit (Takara). Amplified BOT6 fragment was inserted with pGWB402 using pENTR™ Directional TOPO® Cloning Kits (Invitrogen).

### Fungal growth assay

Agar plugs (φ = 5 mm) of ten-day-old fungal cultures grown on Mathurs media were transferred to new Mathurs media plates. Colony formation was determined three days after incubation by measuring the colony radius from center to edge. For measurement of conidia formation, the fungal colonies on Mathurs flasks were suspended with 5 mL of distilled water to obtain conidia. For inoculation on plants, 3 μL of fungal conidia suspension, comprising approximately 5 × 10^3^ conidia/mL, was put onto half MS media containing 50 μM KH_2_PO_4_ (Pi) without sucrose, as described previously^17^. The colony areas were measured by using Image J/Fiji (https://imagej.net/software/fiji/)^51^.

### Fungal inoculation assay

*Arabidopsis thaliana* Col-0, the *abi1-1C* mutant (Umezawa et al., 2009; Col-0 background), and *ein3 eil1* were used in this study. Seeds were surface sterilized with 70% ethanol for 30s, followed by 6% sodium hypochlorite with 0.01% Triton X for 3 min, then washed five times in sterilized water. Sterilized *A. thaliana* seeds were sowed on half MS agarose medium containing 50 µM Pi concentrations. Fungal conidia were obtained from 7-day-old of Ct3, or 5-day-old colonies of Ct4 or transgenic lines formed on Mathurs media. Fungal conidia (3 μL of 5 × 10^3^ conidia/mL) were inoculated 3 cm below the sowed seeds. Plates were placed vertically in a plant growth chamber under a 10:12 hr light: dark cycle at ∼ 22°C (± 1°C) (80 μmol/m^2^s) around 50% humidity. Effects on plant growth by fungal inoculation were evaluated by measuring the fresh shoot weight at 22 or 24 dpi.

### RNA extraction and RNA-seq analysis

To validate the gene expression of Ct4 and BOT6 OX lines (#1, #10, #24) *in vitro*, fungi were incubated at room temperature in 8 mL of liquid Mathur’s media for three days with 100 rpm rotation. The mycelia were collected after removing water as much as possible and frozen in liquid nitrogen. For quantification of gene expression in plant-fungi interactions, 12 dpi or 24 dpi plants were harvested and frozen in liquid nitrogen. These samples were ground to a fine powder by Shake Master NEO (bms). Total RNA was extracted using Nucleospin® RNA (Macherey-Nagel) according to the manufacturer’s instructions. For RNAseq analysis, ∼ 2 µg of RNA samples were sent to Rhelixa for sequencing. After Poly-A selection by NEBNext Poly(A) mRNA Magnetic Isolation Module, the strand-specific libraries were generated by NEBNext Ultra II Directional RNA Library Prep Kit. The generated libraries were sequenced by Illumina NovaSeq 6000 platform (150 bp x 2 paired-end), resulting in approximately 26.7 M reads per sample. The obtained reads were mapped to the Ct4 genome assembly by hisat2 **(version 2.2.1)**^52^. The obtained sam files were converted to the corresponding bam files by samtools **(version 1-19)**^53^. featureCounts **(version 2.0.6)**^54^ were used to obtain the count data of fungal genes from the bam files. In this procedure, the Ct4 gff file was slightly modified based on the predictions derived from the prior RNAseq analysis conducted by Hiruma et al., 2023, focusing on the ABA and BOT genes. Based on the obtained count table, the differentially expressed genes between the WT and each OX line were calculated by the edgeR package **(version 4.0.3)**^55^. The motif analysis was conducted in MEMEsuite **(version 5.5.5)**^56^ (https://meme-suite.org/meme/) using the upstream 500 bp sequences of 1485 fungal genes differentially up-regulated (| log2FC | >1, q < 0.05) in either of OX lines compared with the WT strains. GO analyses were conducted against 1485 fungal genes by Omicsbox platform after Blast2go analysis against all the Ct4 genes **(version 3.1.11)**^57^. Predictions for secreted proteins were conducted by SignalP-6.0 slow-sequential mode^58^. Predictions for candidate effectors were conducted by Effector P-3.0 with default setting^59^. Prediction for secondary metabolism gene clusters was conducted by antismash (https://fungismash.secondarymetabolites.org/)^60^. Predictions for CAZymes were conducted by ddCAN3 (http://www.cazy.org/Home.html)^61^.

### Quantitative real-time PCR

cDNA was synthesized from 100∼200 ng total RNA of mycelia or plant samples, using the PrimeScript RT Master Mix (Takara). We then amplified cDNA in Power SYBR (Thermo) or THUNDERBIRD® Next SYBR™ qPCR Mix (TOYOBO) with 250 nM primers using CFX Opus 384 Real-Time PCR System (BIORAD) in a volume of 10 μL. Primers used in this study are listed in Supplementary Table 6.

### Metabolite analysis

The mycelia of Ct were inoculated into 1 mL of Mathur’s media in a 5 mL test tube. The cultures were incubated at 25°C with a constant rotational speed of 200 rpm for a period of 5 days. Following extraction with 2 mL of acetone, the extracts were evaporated. Subsequently, 1M-HCl was added to the mixture and the resulting mixture was extracted with ethyl acetate. The crude extracts were then dissolved in methanol.

#### For botrydial intermediates

After filtration of the crude extracts, the filtrates were directly analyzed by a GC-MS equipped with a Beta DEX^TM^ 120 fused silica capillary column (0.25 mm × 30 m, 0.25 μm film thickness; SUPELCO) in the following conditions (**Method A**); 100°C for 3 min, 100 – 230°C (rate: 14°C/min), 230°C for 5 min at a flow rate of 1.2 mL/min (helium carrier gas).

#### For ABA intermediates

After filtration of the crude extracts, the filtrates were directly analyzed by a UPLC-MS equipped with a ACQUITY UPLC BEH C18 (2.1 x 50 mm) in the following conditions (**Method B**); 0–1 min = 10%B, 1–3 min = 10–95% B, 3-5min = 95% B (A: H_2_O + 0.1% of formic acid, B: CH_3_CN + 0.1% of formic acid) at a flow rate of 0.7 mL/min; column temp. 40°C.

#### Profiling fungal metabolites during solid medium incubation

Mycelia of Ct were inoculated into a solid medium containing polished rice (1 g) in 5 mL test tube. The culture was incubated at 25°C for 5 days. After extraction with Ethyl acetate, the extract was concentrated in vacuo to afford crude extracts.

#### For botrydial intermediates

After filtration of the crude extracts, the filtrates were directly analyzed by a GC-MS in **Method A**.

#### For abscisic acid intermediates

The crude extracts were evaporated and the residues were then dissolved in MeOH. After filtration, the filtrates were directly analyzed by a UPLC-MS in **Method B**.

#### For new sesquiterpenes

After filtration of the crude extracts, the filtrates were directly analyzed by a GC-MS in **Method A.**

### Microscopy

Fungal hyphae infected on *A. thaliana* roots (16 dpi), or leaves (22 dpi) were visualized by confocal microscopy (OLYMPUS FLUOVIEW FV3000) using WGA-FITC (Sigma). A 20.0 × UPLXAPO20X objective lens (0.8 numerical aperture) was used. Excitation dichroic mirrors DM405/488 and variable barrier filters (VBF) of 487 nm–545 nm were used for the detection of fluorescent signals. For lectin staining, *A. thaliana* roots were incubated overnight with 80% methanol. The roots were then transferred to chloral hydrate (2.5 g/mL in water) and incubated overnight. The roots were stained with wheat germ agglutinin (WGA) conjugated to a FITC solution (5 μg/ml; Sigma) for 2 hr at room temperature after washing with PBS buffer.

### Dual-Luciferase assay

To assess the binding activity of CtBOT6 to the promoter sequences of *CtABA3* and GENE_00001199, *Nicotiana benthamiana* was used to transiently express CtBOT6 protein and the promoters of interest. Each vector was transformed into *Agrobacterium* strain GV3101 by electroporation. For infiltration, Agrobacterium overnight cultured cells were harvested by centrifugation and then resuspended in MMA induction buffer (1 L of MMA: 5 g of Murashige and Skoog salts (FUJIFILM Wako Pure Chemicals), 1.95 g of MES, 20 g of sucrose, and 200 μM acetosyringone, pH 5.6). The suspensions were adjusted to OD600 = 0.2 and mixed the Agro carrying reporter with the Agro carrying the CtBOT6 construct at 1:5 ratio, and then infiltrated into *N. benthamiana* leaves. The leaves at 3 dpi were tested for the reporter assay. Firefly (*Photinus pyralis*) luciferase was used as a reporter, and *Renilla* (*Renilla reniformis*) luciferase was used as a control. The activities of luciferase were measured using Dual-Luciferase® Reporter Assay System (Promega), according to the manufacturer’s instructions. The Luminescence was monitored using Tristar 3 (BERTHOLD).

### Statistical analysis

For quantitative PCR, statistical analysis was performed in R using the linear mixed model function (lmer) from the package lme4 following a method described in Mine *et al*., 2017. The following models were fit to the relative Ct value data compared to *plantACTIN* or *CtACTIN*: Ctyre = Yy + Rr + eyr or Ctgyre = GYgy + Rr + egyr, where GY, genotype:treatment interaction and random factors; R, biological replicate; e, residual. Differences between estimated means were compared in twoltailed *t*ltests. For the *t*ltests, the standard errors appropriate for the comparisons were calculated with the variance and covariance values obtained from the model fittings. The Benjamini-Hochberg method was applied to correct for multiple hypothesis testing when all pairwise comparisons of the mean estimates were made. For calculating Cohen’s d effect size, the R package effsize was used. For Pearson’s product-moment correlation, log2-transformed Shoot fresh weight and Ct (*plant ACTIN*)- Ct (*CtBOT6*) value were used for calculation.

## Additional information

Raw sequencing data of RNA-seq transcriptomic data have been deposited in DDBJ (PRJDB17668 (PSUB022574)).

## Acknowledgments

We thank Sachiko Minezaki for technical assistance. We thank Yasuyuki Kubo for the published materials. We thank Akira Mine for the unpublished material (pAT006-Fluc-iG5_Rluc-iFAD2_F vector). This work was supported in part by the JSPS KAKENHI Grant (JP23H02210, JP22H02204 (A.M.)), the JST grant (JPMJAN23D4, JPMJFR200A), JST SPRING (JPMJSP2108), and The Uehara Memorial Foundation (A.M.).

## Author contributions

KH initiated the project. RU and KH directed the research. RU, JT, MN, and HH conducted the experiments. RU and KH conducted RNA-seq analyses. RU, JT, AM, and KH analyzed the data. RU and KH wrote the manuscript with feedback from all authors.

---

**Extended Data Figure 1.**
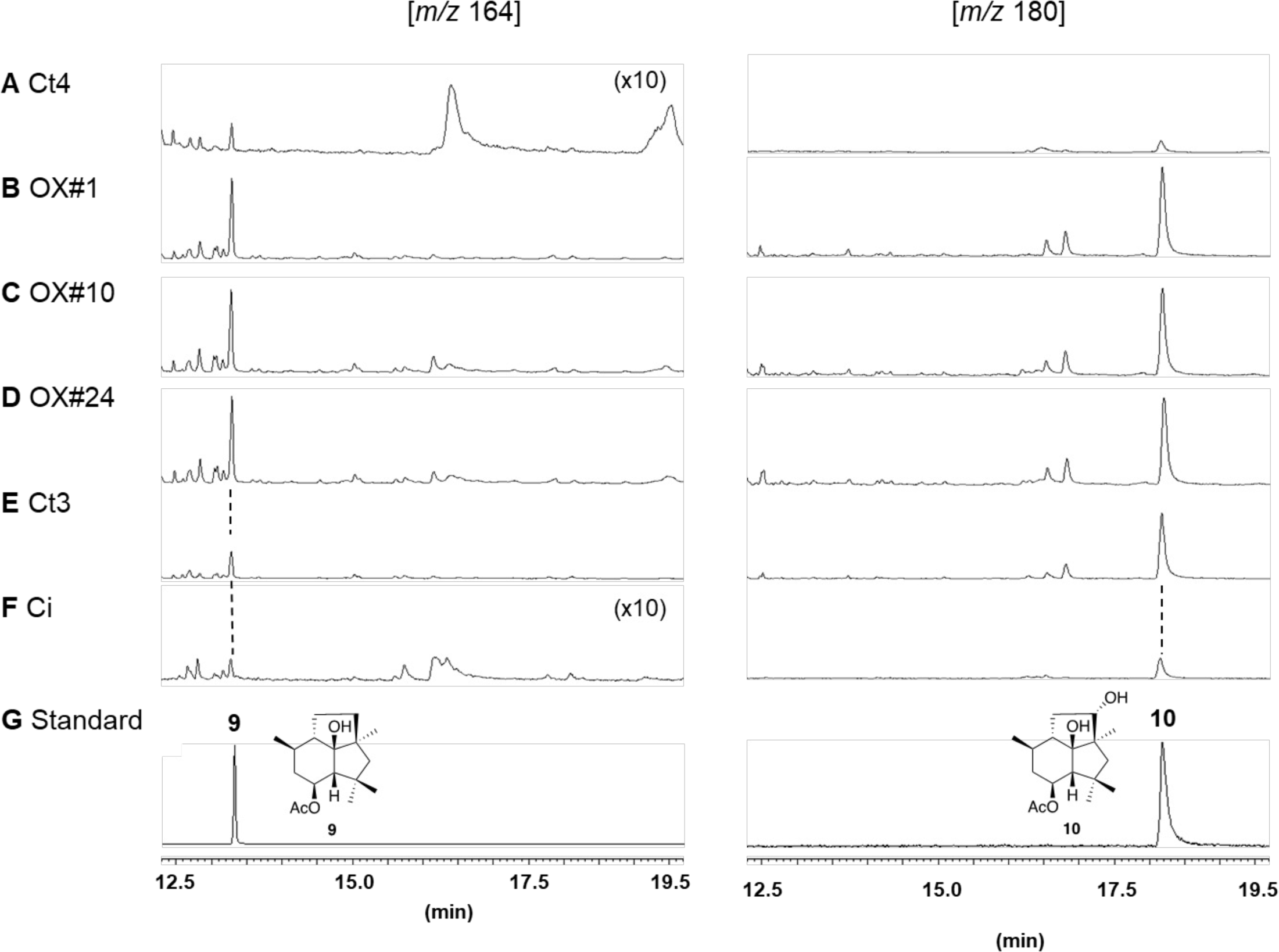
Constitutive expression of *CtBOT6* increases the production of the biosynthetic intermediates of botrydial in rice media. GC-MS profiles: The metabolites (rice media) from (A) Ct4, (B) OX#1, (C) OX#10, (D) OX#24, (E) Ct3, and (F) Ci. (G) standard compounds.

**Extended Data Figure 2.**
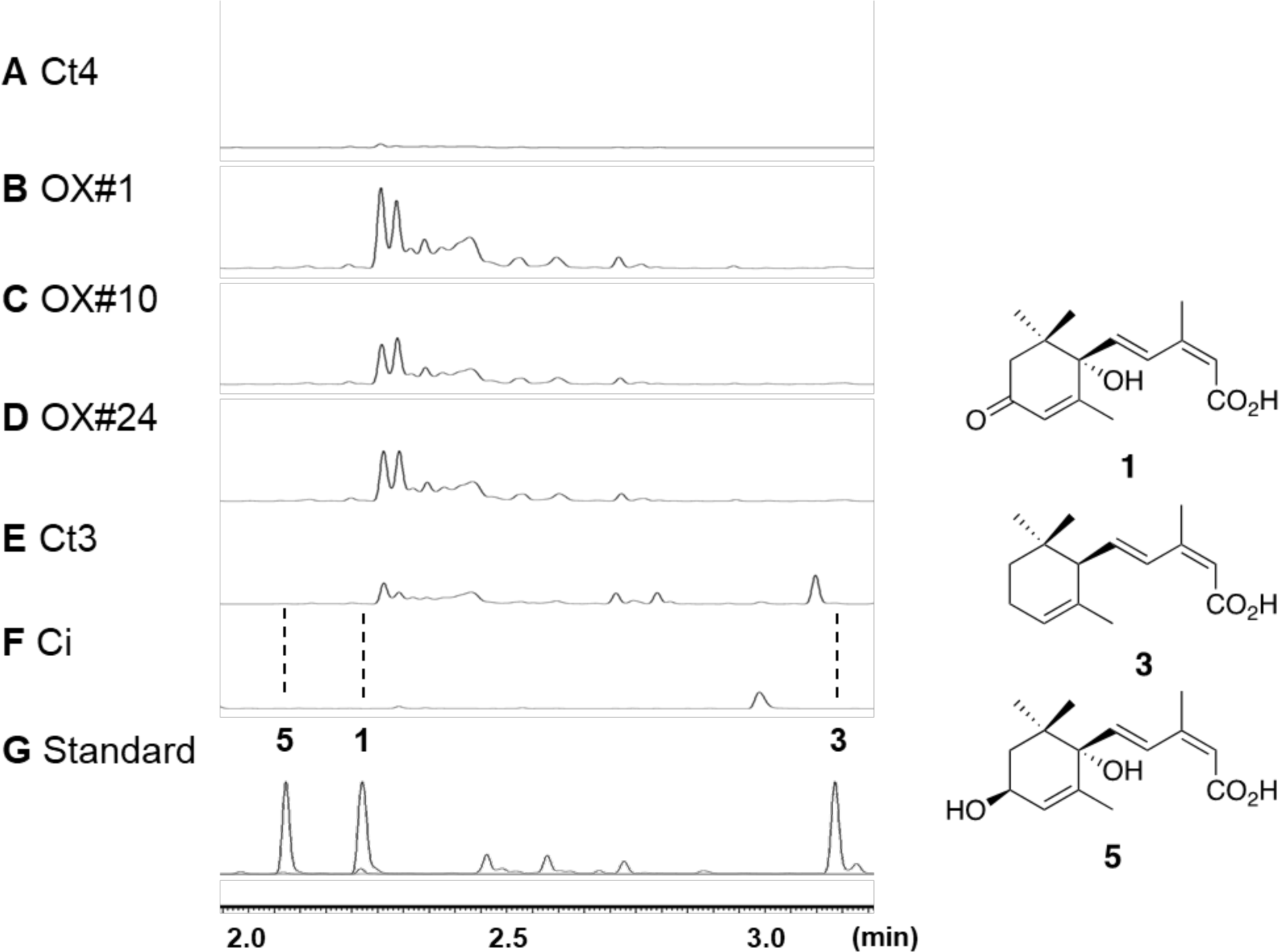
Measurements of ABA and the intermediate metabolites in rice media. UPLC profiles (264 nm): The metabolites (rice media) from (A) Ct4, (B) OX#1, (C) OX#10, (D) OX#24, (E) Ct3, and (F) Ci. (G) standard compounds.

**Extended Data Figure 3.**
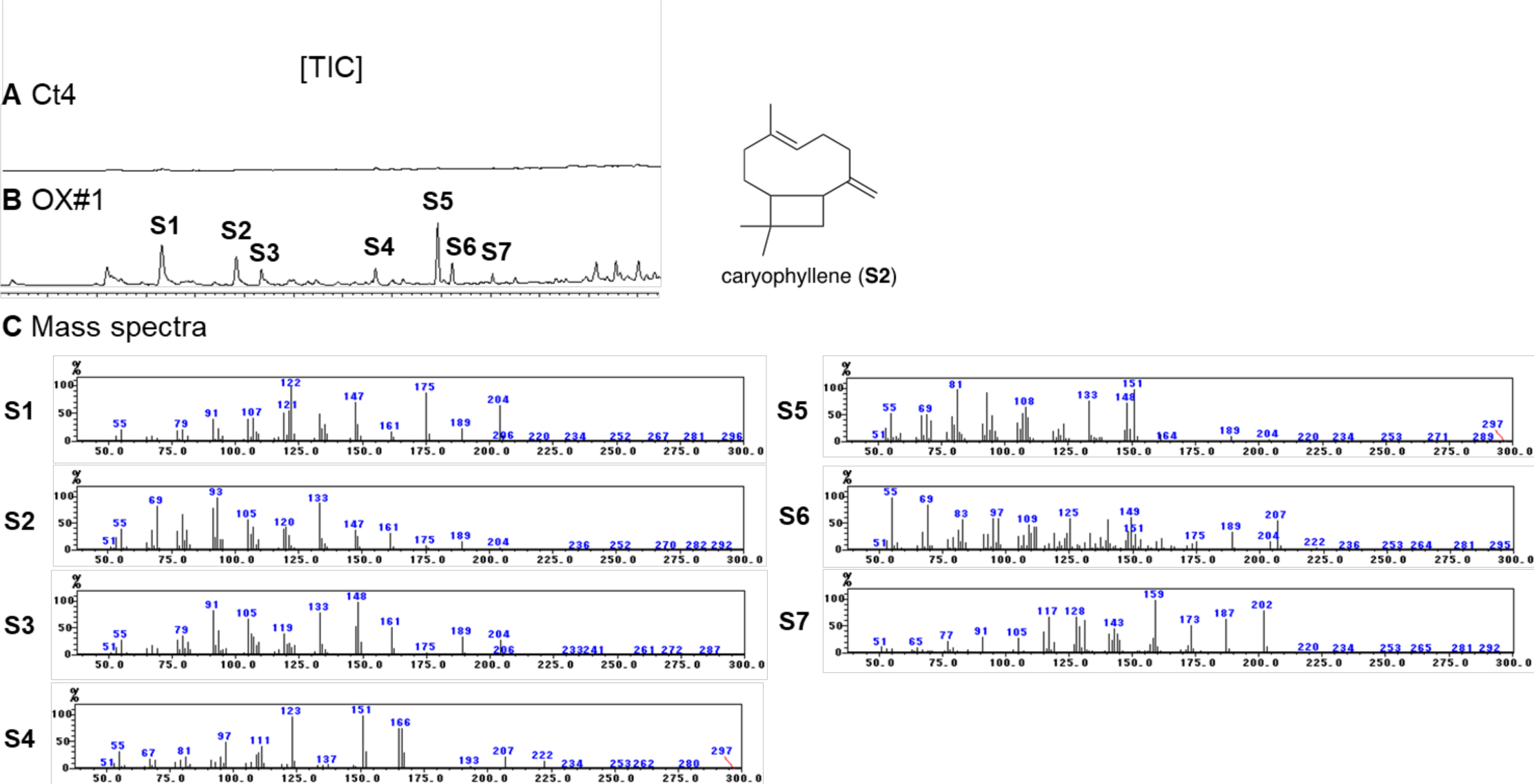
Production of new metabolites in OX#1. GC-MS analysis of the metabolites (rice media) from (A) Ct4 and (B) OX#1. (C) Mass spectra of new metabolites S1-S7. Among them, S2 was proposed to be caryophyllene based on the similarity analysis^62^.

**Extended Data Figure 4.**
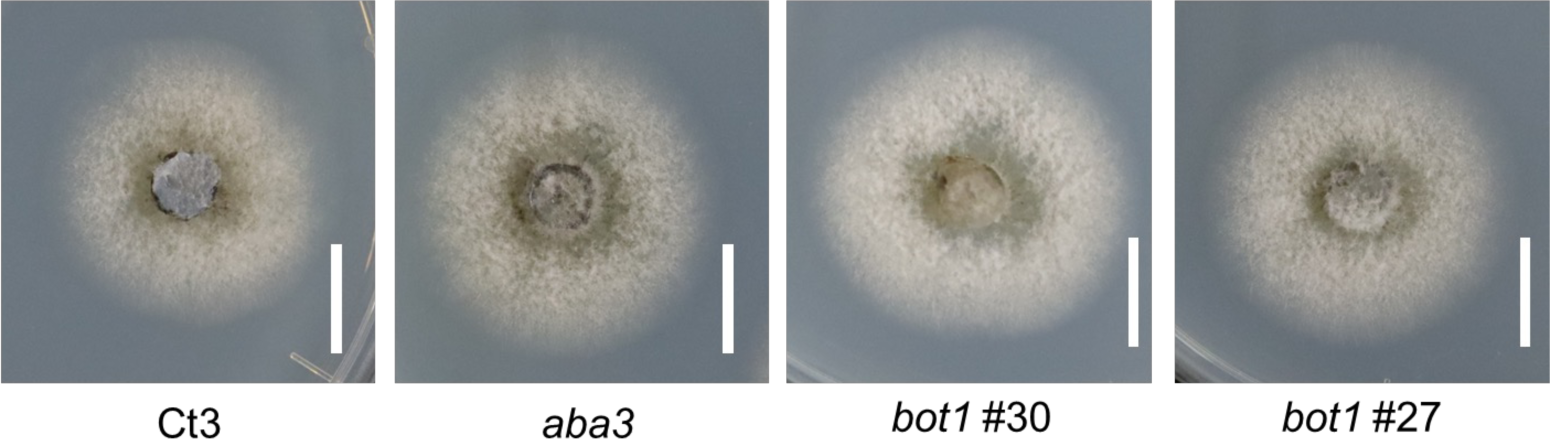
Gene disruption of *CtABA3* and/or *CtBOT1* do not alter appearance of fungal colony *in vitro* Growth of Ct3 WT and Ct3 *aba3* and *bot1* (#30 and #27) mutants in Mathur’s nutrient media after 3 days of incubation. Bars = 1 cm.

**Extended Data Figure 5.**
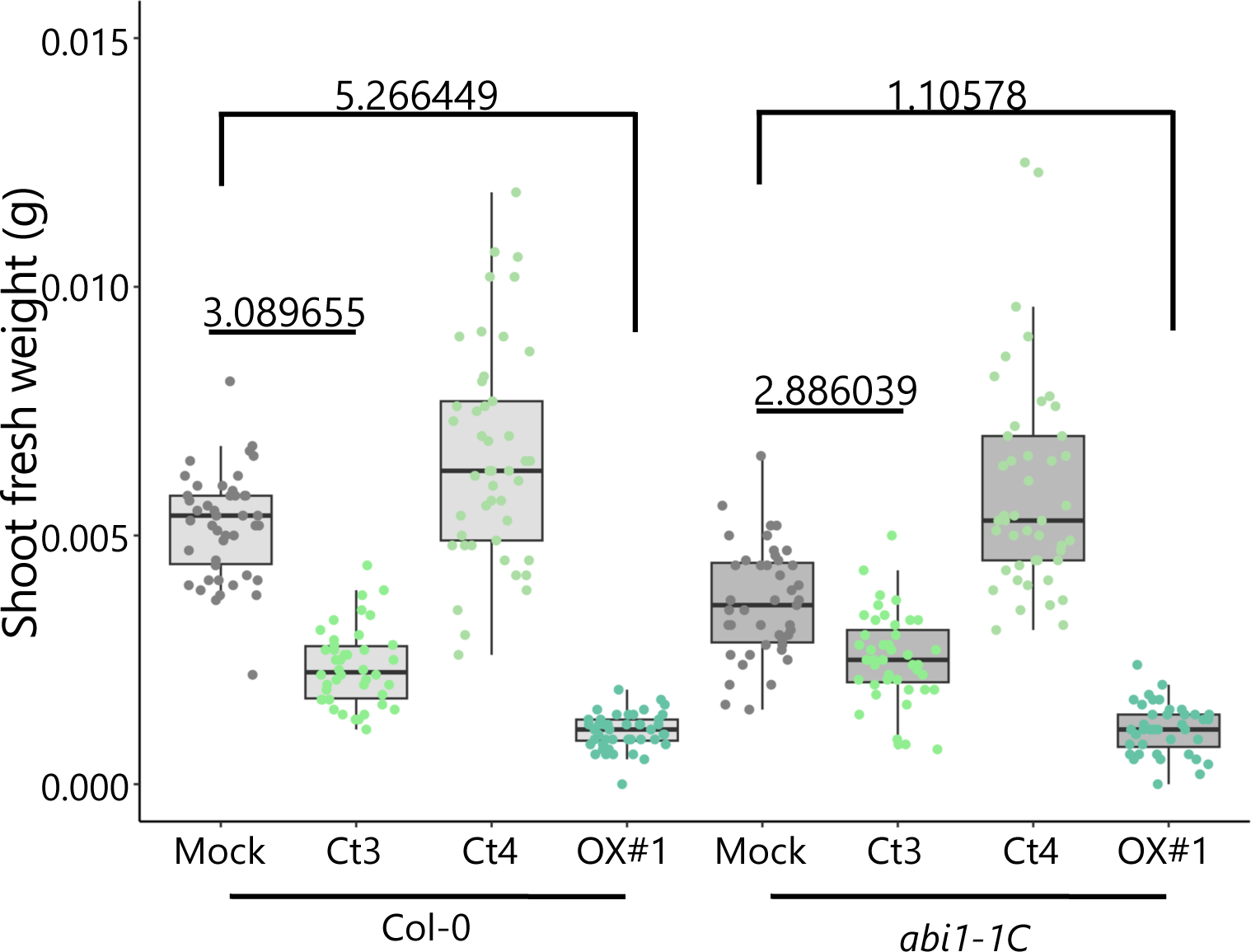
Quantitative data of *A. thaliana* shoot fresh weight in Pi-limiting conditions under fungal infection. Seedlings were harvested at 24 dpi. Each dot represents individual plant samples (For Col-0, Mock: *n* = 46, Ct3: *n* = 42, Ct4: *n* = 45, OX#1: *n* = 44; for *abi1-1C*, Mock: *n* = 43, Ct3: *n* = 43, Ct4: *n* = 45, OX#1: *n* = 40 biologically independent samples). Numerals indicate effect size. A larger effect size designates a stronger relationship between two variables.

**Extended Data Figure 6.**
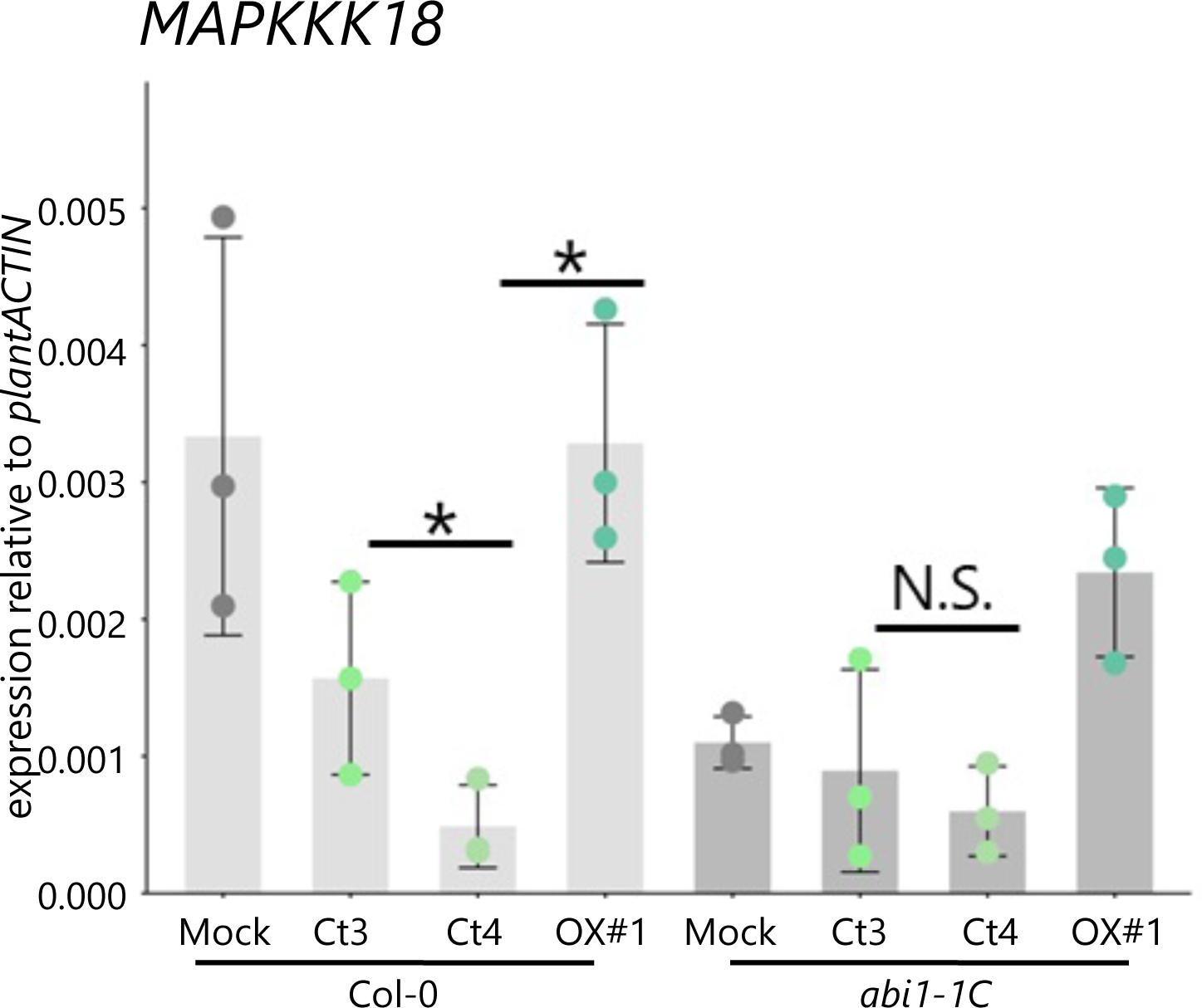
Plant ABA signaling responses under fungal infection Plant gene expressions in roots at 12 dpi were measured by RT-qPCR and normalized with plant *ACTIN*. Error bars indicate ±SD. *n* = 3 biologically independent samples from three independent experiments. The asterisks indicate a significant difference (two tailed *t*-test, *p* = 0.01091 in Col-0 Ct3 v.s. Col-0 Ct4, *p*= 0.00025 in Col-0 Ct4 v.s. Col-0 OX#1). N.S. means no significant difference.

**Extended Data Figure 7.**
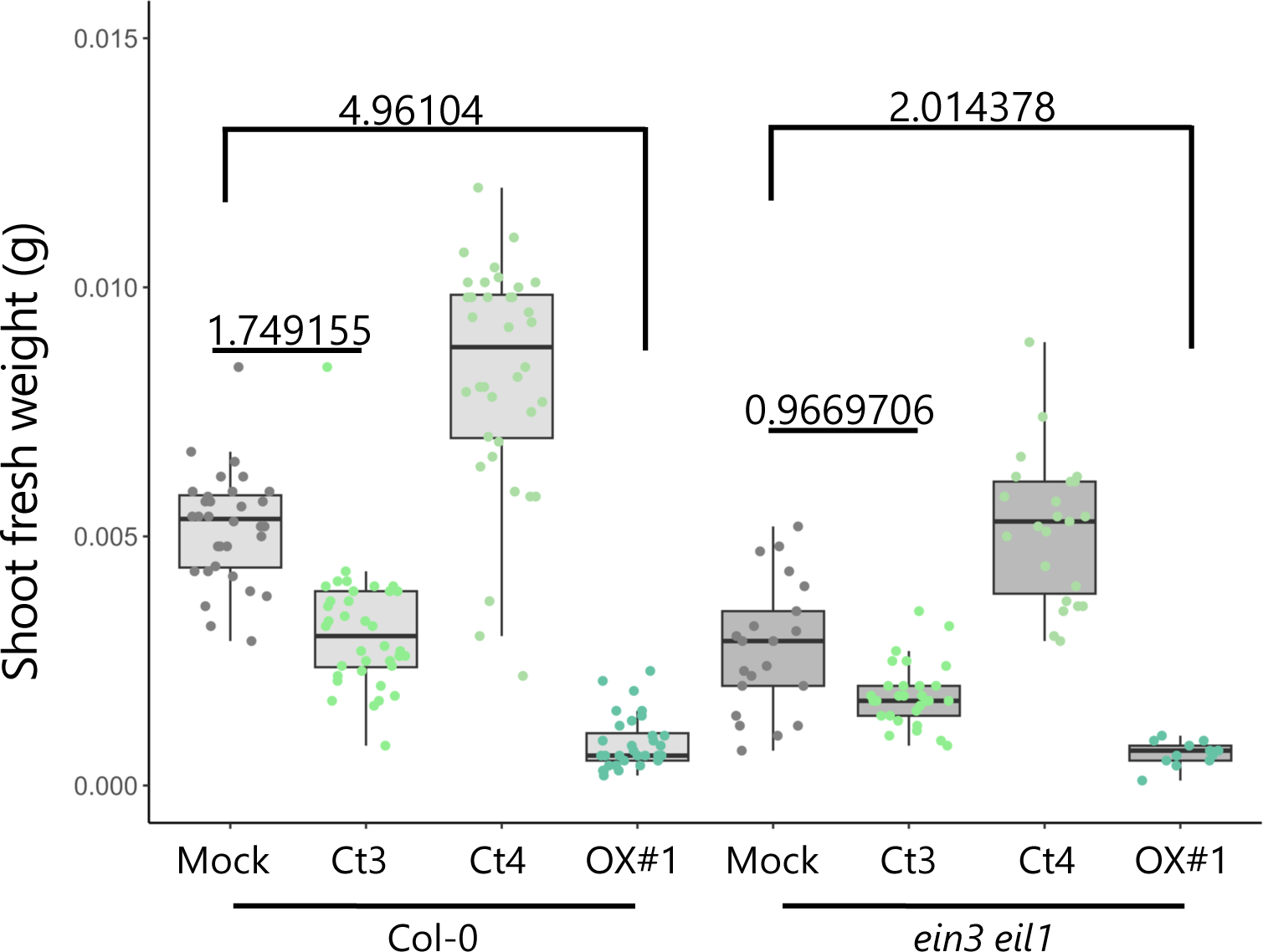
Quantitative data of *A. thaliana* shoot fresh weight in Pi-limiting conditions under fungal infection. Seedlings were harvested at 24 dpi. Each dot represents individual plant samples (For Col-0, Mock: *n* = 32, Ct3: *n* = 36, Ct4: *n* = 36, OX#1: *n* = 32; for *ein3 eil1*, Mock: *n* = 21, Ct3: *n* = 29, Ct4: *n* = 23, OX#1: *n* = 13 biologically independent samples). Numerals indicate effect size. A larger effect size designates a stronger relationship between two variables.

**Extended Data Figure 8.**
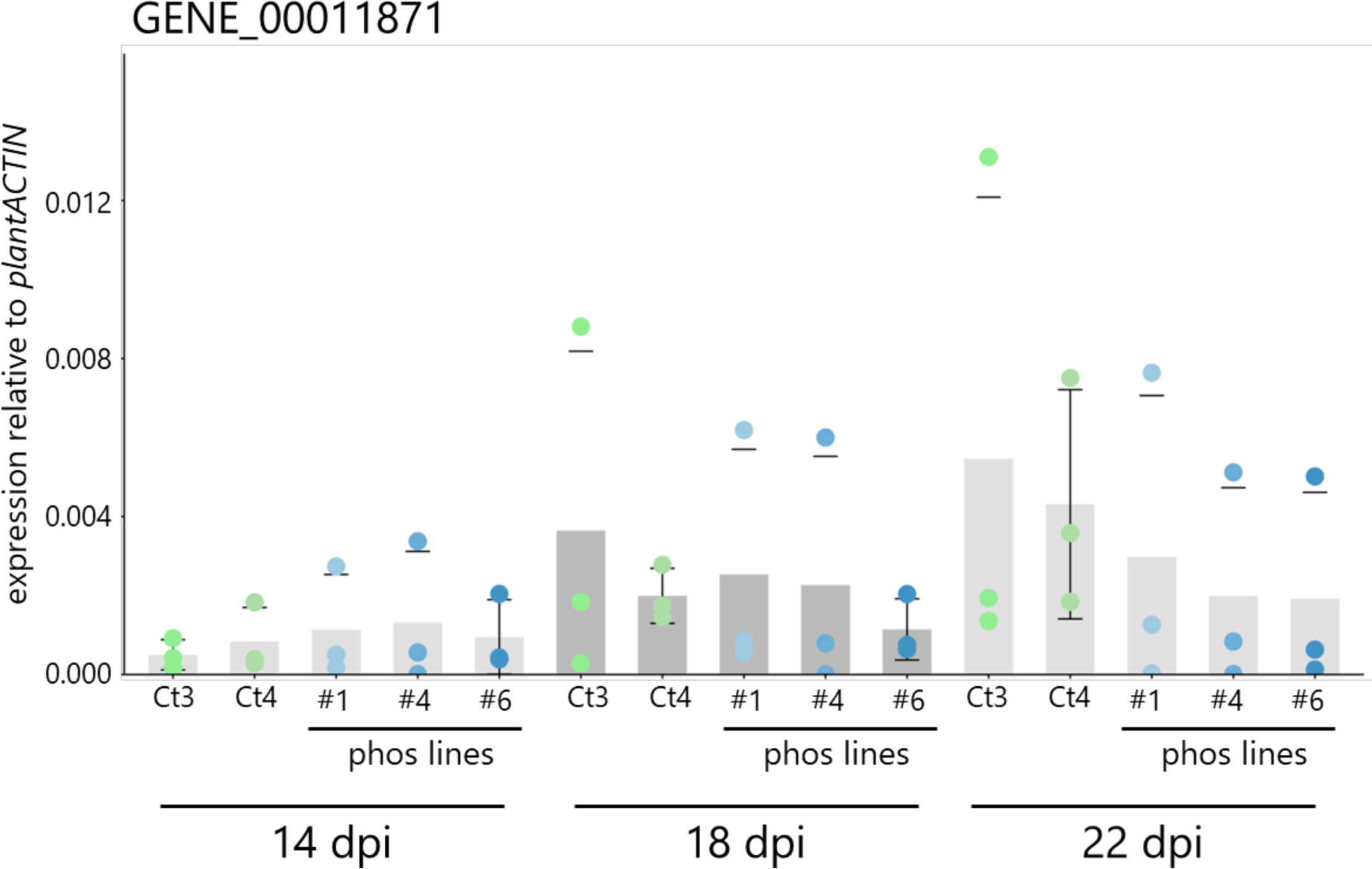
The expression dynamics of fungal phosphate transporter (GENE_00011871). Fungal gene expressions in roots at 14, 18, and 22 dpi were measured by RT-qPCR and normalized with plant *ACTIN*. Error bars indicate ±SD. *n* = 3 biologically independent samples from three independent experiments. Results from two technical replicates were combined.

